# Functional integration of the bacteriophage T4 DNA replication complex: The multiple roles of the ssDNA binding protein (gp32)

**DOI:** 10.1101/2025.09.05.674579

**Authors:** Claire S. Albrecht, Brett Israels, Jack Maurer, Peter H. von Hippel, Andrew H. Marcus

## Abstract

Single-stranded DNA binding protein (gp32) serves as the central regulatory component of the multi-subunit T4 bacteriophage DNA replication system by coordinating the system’s three functional sub-assemblies, resulting in phage DNA synthesis in T4-infected *E. coli* cells at the high speeds (∼1,000 nts s^-1^) and the high fidelity (< 1 error per 10^7^ nts) required for genomic function within this cellular eco-system. Gp32 proteins continuously bind to, slide as cooperatively-linked clusters on, and un-bind from transiently exposed single-stranded (ss) DNA templates to carry out their coordinating functions, as well as to protect genomic sequences from nuclease activity and block the formation of interfering secondary structures. The N-terminal domains (NTDs) of gp32 mediate cooperative interactions within ssb clusters, but the roles of the disordered C-terminal domains (CTD) in the nucleation of gp32-ssDNA filaments at ss-dsDNA junctions are less well understood. We here present microsecond-resolved single-molecule Förster resonance energy transfer studies of the initial steps of gp32 assembly on short oligo-deoxythymidine lattices of varying lattice length and polarity near model ss-dsDNA junctions. These data are analyzed to define the molecular steps and related free energy surfaces involved in initiating gp32 cluster formation, which show that the nucleation mechanisms and regulatory interactions driven by gp32 proteins at ss-dsDNA junctions are significantly directed by lattice polarity. We propose a model for the role of the CTDs in orienting gp32 monomers at lattice positions close to ss-dsDNA junctions that suggests how intrinsically disordered CTD domains might facilitate and control non-base-sequence-specific binding in both the nucleation and the dissociation of the gp32-ssDNA filaments involved in phage DNA replication and related processes.

## I. Introduction

The DNA replication system of bacteriophage T4 serves as a simple model for how higher organisms (including humans) control genome function (1, 2). Three separate protein sub-assemblies collaborate to open the DNA duplex and expose the single-stranded DNA templates, which are then each ‘copied’ by DNA polymerase to synthesize new ‘daughter’ DNA strands from nucleotide precursors. The polymerase molecules and the DNA strands being copied and extended are held together during the synthesis process by multimeric ‘sliding clamps,’ which encircle the polymerase-ssDNA complexes and prevent them from dissociating after each synthesis step. The three sub-assembly complexes are controlled and integrated by the single-stranded DNA binding (ssb) protein of T4 (called gene product 32, or gp32, accession ID: PO3695), which serves as the central regulatory component of the system by binding at the replication fork and primer-template junctions to coordinate the initiation of DNA replication, transcription and repair (3, 4).

During DNA synthesis, short clusters of ssb proteins bind to, unbind from, and slide on exposed ssDNA templates to initiate and integrate the enzymatic cycles of the three spatially separated replication complex sub-assemblies (*i*.*e*., the helicase-primosome, the DNA polymerases and the clamp-clamp loader complex) (5). Cooperative interactions between adjacent ssb proteins bound to ssDNA (6, 7) result in the initiation and assembly of structured nucleoprotein filaments, which protect the exposed ssDNA templates from degradation by nucleases and prevent the formation of secondary structures that would interfere with and slow the replication process (8-10). In addition, ssb protein-DNA filaments align the template strands as they are fed into the replicative polymerases and out from the primosome complex at the replication fork junction (11-14). Such direct interactions between ssbs and the replication protein sub-assemblies work to integrate leading and lagging strand DNA synthesis (15).

The gp32 protein has three functional domains: a core DNA binding domain (DBD), an acidic C-terminal domain (CTD) and a basic N-terminal domain (NTD). The core domain contains a highly electropositive binding cleft, which interacts with the negatively charged sugar-phosphate backbone of ssDNA (6, 8, 12, 16, 17), while the CTD is a disordered, negatively charged polypeptide tail that is responsible for protein-protein interactions between gp32 and other protein sub-assemblies (4, 11, 18-21). The NTD functions to mediate cooperative interactions between adjacent gp32 molecules bound to ssDNA lattices (4, 6, 7, 20). In Fig. 1*A*, we schematically depict three conformations of the gp32 protein that function during different stages of DNA replication. When gp32 is not bound to ssDNA, the CTD can occupy a conformation in which the binding cleft is partially occluded. This creates competition between the CTD and ssDNA at the binding cleft, which competition is mediated by conformational changes of the CTD (18). The gp32 protein can also take up conformations in which the CTD is bound to the ‘core’ DNA binding domain (DBD), or in which the CTD is extended away from the DBD.

**Figure 1.**
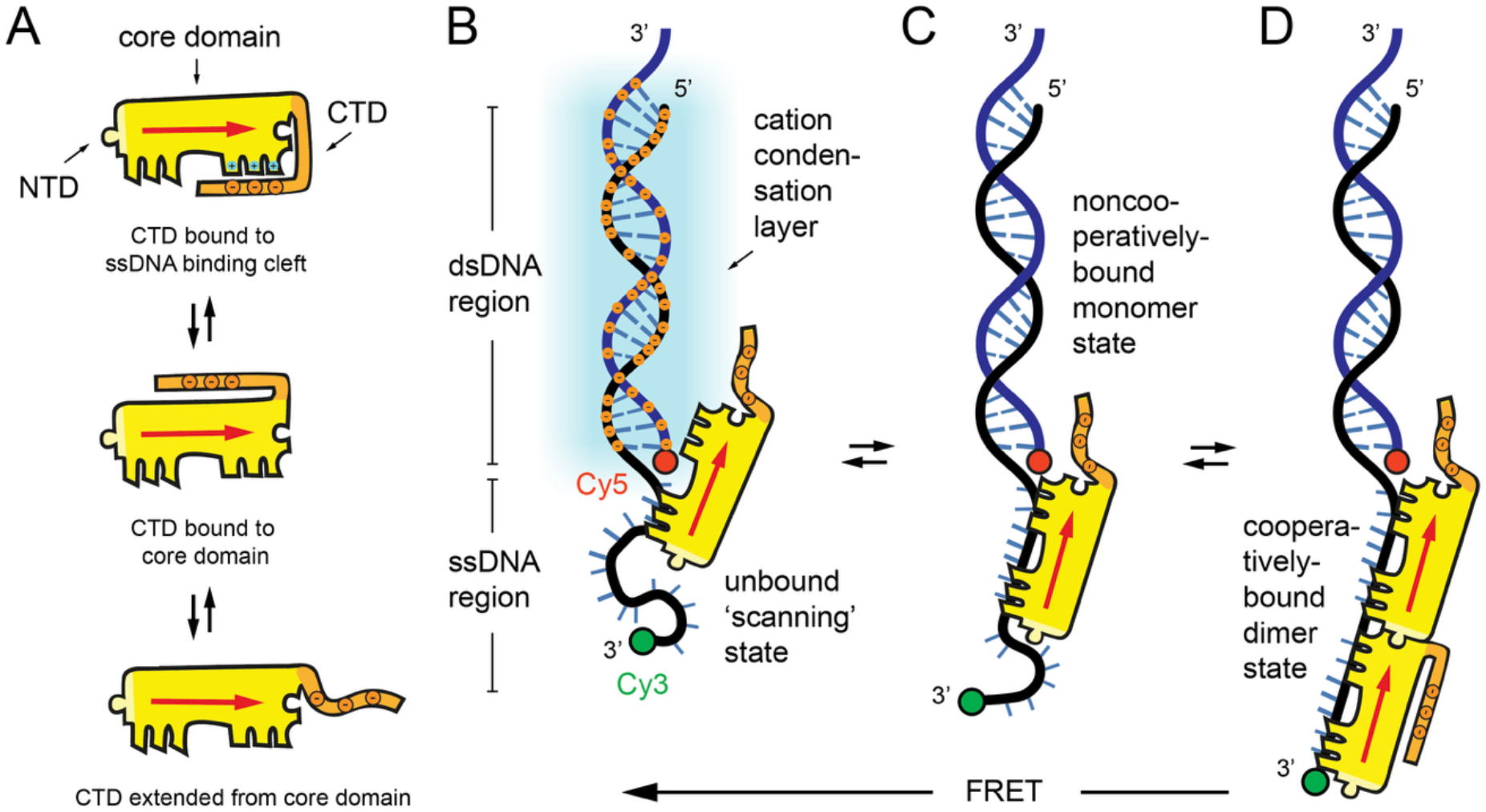
(***A***) Schematic of the gp32 protein illustrating the N-terminal domain (NTD), the core ssDNA binding domain (DBD) and the disordered C-terminal domain (CTD). The negatively charged CTD can be bound to the positively charged binding cleft (Panel A – top), to the core domain (Panel A – middle), or extended away from the core domain (Panel A – bottom). The core domain contains a ssDNA binding site of length *s* = 7 nts. Note the directional red arrow points from the NTD toward the CTD. (***B***) The duplex region of the ss-dsDNA junction is surrounded by a net positive ion cloud (indicated by blue circles), which attracts and orients the CTD of the gp32 protein near the junction. The extension of the ssDNA region (14, 15 nts) is monitored using a Cy3/Cy5 donor-acceptor FRET pair, as shown. (***C***) A non-cooperatively bound gp32 monomer near the junction is a ‘productive’ intermediate on the pathway to form (***D***) a cooperatively bound dimer cluster. The states illustrated in panels B – D exhibit successively decreasing FRET values.

Gp32 binds initially to ssDNA as a monomer at ‘physiologically equivalent’ salt concentrations (22). Our current *in vitro* studies were carried out at salt concentrations ([NaCl] = 100 mM and [MgCl_2_] = 6 mM) that result in protein-DNA binding constants that are the same as those measured in living *E. coli* cells (23). The ‘binding site size’ (functional length) of the gp32 binding cleft (*s*) is ∼7 nucleotide residues (nts) (3, 16, 24). The initial binding of gp32 monomers to ssDNA results in their random distribution along the exposed ssDNA lattice sequence, with a binding preference for pyrimidine-rich sequences in which the stacking free energies are relatively small and therefore provide higher affinity binding sites than those containing significant quantities of more highly stacked purine residues that must be unstacked to permit cooperative gp32 binding (20, 25). These initially separate gp32 monomers then translocate along the ssDNA sequences to form cooperatively bound ssb clusters that can melt highly AT-rich regions of dsDNA (3). Interactions between ssb monomers and double-stranded (ds) DNA are non-cooperative and ∼10^4^ times weaker than the cooperative interactions that form between ssb monomers and ssDNA (3). On ssDNA lattices of sufficient length to bind multiple proteins, gp32 can form cooperatively bound (ssb)_*n*_-ssDNA ‘clusters.’ The cooperativity factor between adjacent gp32 monomers bound to ssDNA is ∼10^3^ (26, 27), and contiguously bound (and relatively short) gp32 clusters can undergo one-dimensional ‘sliding’ along ssDNA lattice (4, 20, 27-29).

Gp32 proteins bind in a preferred direction along the ssDNA lattice due to the asymmetry of the protein-ssDNA interaction. However, little structural information is available about the polarity preference of gp32-ssDNA filaments, or how this preference is affected by the presence of other DNA-bound proteins or nearby secondary structures such as ss-dsDNA junctions. In prior work, fluorescent base analogue-substituted DNA constructs were used to ‘map’ the local conformations of nucleobases within the binding cleft of gp32 proteins (16, 17). These studies showed that: (*i*) the gp32 protein monomer has two tight binding interactions at positions 5 and 6 within the binding cleft in the 5’ → 3’ direction of the ssDNA lattice (16), and (*ii*) cooperatively bound clusters tend to ‘slide’ towards the 5’ end of the ssDNA lattice (20). By combining these results with structural information about the gp32 core domain (8) it was inferred that the protein binds to the ssDNA lattice with its CTD oriented preferentially toward the 5’-end of the lattice and its NTD toward the 3’-end.

However, we also note that recent crystal structures of gp32 in complexation with the repair helicase Dda and model ss-dsDNA fork constructs suggest the opposite polarity preference for that complex (30). These findings suggest that the polarity preference of gp32 proteins bound to ssDNA lattices may, in addition to strand polarity, also be regulated by the presence of other proteins and protein sub-assemblies. We thus expect gp32 binding and function near ss-dsDNA junctions to depend on the details of the system under study and understanding it better may provide new insights into how ssb proteins mediate their interactions with replication protein sub-assemblies and with one another.

In the work presented in this paper, we summarize *microsecond-resolved* single-molecule Förster resonance energy transfer (smFRET) studies of gp32 dimer assembly on short oligo-deoxythymidine [oligo(dT)_*n*_] lattices near model ss-dsDNA ‘primer-template’ (p/t) DNA junctions. These experiments serve to define the initiation phase of gp32 binding to ssDNA lattice sites located immediately adjacent to replication forks and p/t junctions and permit us to investigate the sensitivity of the (gp32)_2_-oligo(dT)_*n*_-dsDNA assembly pathway to ssDNA oligo(dT)_*n*_ lattice lengths (*n* = 14 and 15 nts) and polarities (3’ and 5’) relative to the oligo(dT)_*n*_-dsDNA junction. In each of our oligo(dT)_*n*_-dsDNA constructs (see Table 1, below), the distal end of the overhanging oligo(dT)_*n*_ chain is labeled with FRET-donor chromophore Cy3, and the conjugate DNA strand labeled at the ss-dsDNA junction with FRET-acceptor chromophore Cy5. We note that these same constructs were used in previous smFRET / gp32 assembly experiments from our laboratories (24, 31). However, those earlier studies did not have the microsecond temporal resolution of the current work, and thus the kinetic considerations could not be used to facilitate the identification of structural intermediates in the initial assembly of the various DNA construct-gp32 complexes.

**Table 1.**
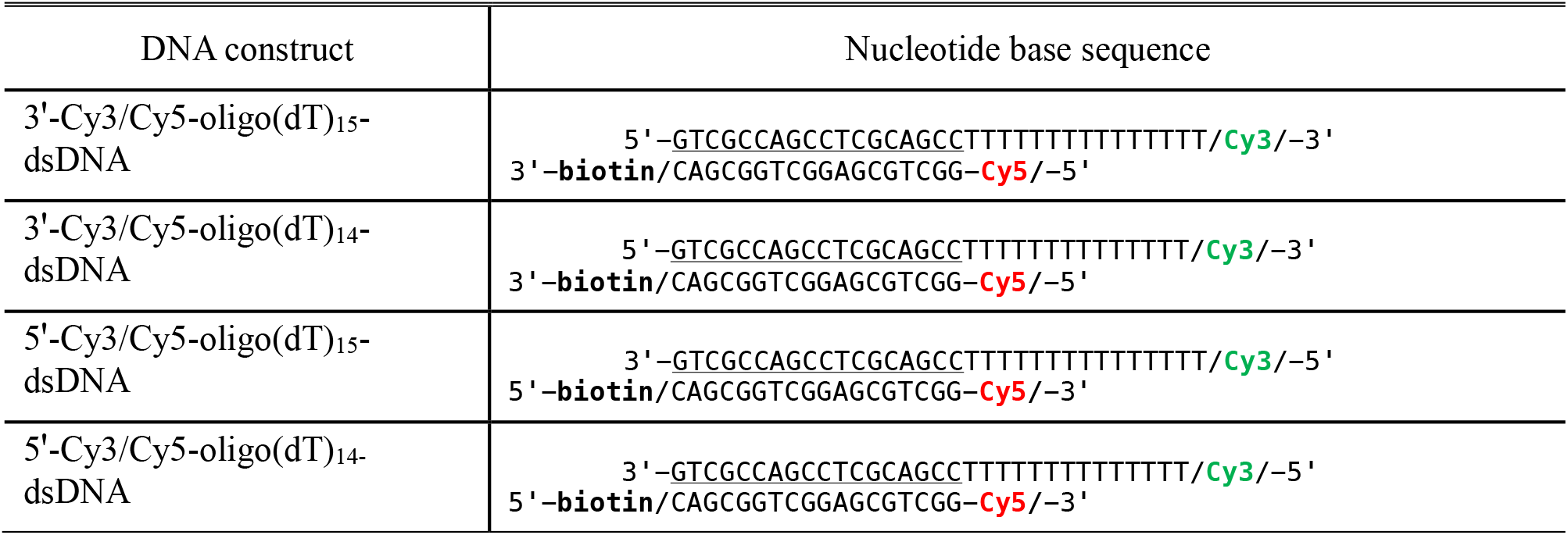
Nucleotide base sequences for ss-dsDNA constructs studied in this work. Horizontal lines indicate regions of complementary base pairing within the duplex regions of the constructs.

For the DNA constructs that we studied there was precisely enough room on the *n* = 14 nts ssDNA segment to permit the cooperative assembly of a single gp32 dimer cluster, while the *n* = 15 nts constructs provided two possible positions for the cooperatively-bound gp32 dimer state, spanning either nt positions 1 – 14 or positions 2 – 15. Our earlier studies (24) showed that gp32 proteins bind to, and dissociate from, the ss-dsDNA junction as monomers on a tens-of-milliseconds time scale. Moreover, the pathway to form a cooperatively-bound dimer cluster includes non-cooperatively bound monomer intermediates in which the monomer must be positioned either at the oligo(dT)_14,15_-dsDNA junction or near the distal end of the oligo(dT)_14,15_ sequence to permit the binding of a second gp32 protein with room to form a properly aligned oligo(dT)_14,15_-dsDNA-(gp32)_2_ DNA replication initiation cluster.

The mechanism by which the gp32 monomer selects its preferred orientation and position relative to the oligo(dT)_14,15_-dsDNA junction in the initial nucleation-assembly step remains an unresolved issue. In Figs. 1*B* – 1*D*, we depict a hypothetical mechanism for the non-cooperative monomer binding of gp32 at the p/t oligo(dT)_14,15_-dsDNA junction. In the absence of bound protein, the oligo(dT)_14,15_ sequences of the DNA constructs rapidly interconvert between a discrete set of conformational macrostates in which most of the flanking bases of the potential gp32 monomer binding sites are unstacked (32). The presence of a few stacked flanking bases at interspersed positions along the chain results in conformational macrostates with more compact end-to-end distances. At positions near the ss-dsDNA junction, the duplex region is partially stabilized by a condensation layer of monovalent sodium ions that screen the repulsive interactions between inter-strand phosphate residues within the sugar-phosphate backbones. Figure 1*B* illustrates a gp32 monomer with its CTD extended towards the duplex region of the ss-dsDNA junction. The gp32 monomer is stabilized and oriented near the ss-dsDNA junction due to the presence of the cation condensation layer, which interacts favorably with the negatively charged CTD. Such an unbound ‘scanning’ state of the CTD of the gp32 monomer might well serve as a precursor to the positioning of a non-cooperatively bound monomer near the oligo(dT)_14,15_-dsDNA junction (Fig. 1*C*), which then could serve as an intermediate on the assembly pathway of a cooperatively-bound dimer gp32 cluster (Fig. 1*D*).

In the following sections of this paper, we show that the assembly mechanisms depicted in Figs. 1*B* – 1*D* are consistent with the results of our smFRET studies of oligo(dT)_*n*_ lattice fluctuations near p/t DNA junctions in the presence of gp32 proteins. Our findings provide new insights into the roles played by the ss-dsDNA junction and the gp32 CTD in the nucleation and association steps of gp32 binding and assembly that are involved in the control of the initiation stages of the overall DNA replication process.

## II. Materials and Methods

### Sample preparation and model p/t-DNA constructs

Model p/t-DNA constructs labeled with Cyanine 3 (Cy3) / Cyanine 5 (Cy5) donor-acceptor FRET chromophore pairs were purchased from Integrated DNA Technologies. The base sequences of these constructs are shown in Table 1. The gp32 protein (UniProt SSB_BPT4: P03965) was prepared and purified as described in (31). Additional details of the sample chamber construction, cleaning procedures and sample preparation are included in the SI, which are essentially the same as reported in prior work (24, 31, 32).

### Microsecond-resolved smFRET and kinetic network model analysis

smFRET experiments were carried out at 1-μs resolution using instrumentation previously developed in our laboratory (24, 32). We analyzed our data using the kinetic network model to simulate experimental time-correlation functions (TCFs) and probability distribution functions (PDFs), as described in previous work (24, 32-34). Additional experimental details and theoretical considerations specific to this work are provided in the SI.

## III. Results

To understand how these processes proceed in detail, we performed microsecond-resolved smFRET experiments on four ss-dsDNA constructs in the presence of gp32 (ssb) protein (32). These constructs all contained an oligo(dT)_*n*_ overhanging ‘tail’ region that varied in length (*n* = 14 or 15 nts) and polarity (3’ or 5’ orientations). The sequences used for each of the four constructs are shown in Table 1. The constructs were fluorescently labeled using the Cy3/Cy5 donor-acceptor FRET pair, as also indicated in Table 1. Both cyanine dyes were attached to the DNA framework using flexible linkers as described in previous papers (32, 35).

In one series of experiments, presented in detail here in the Results section as an example of our approach, we examine the assembly pathway of the gp32 dimer onto the 3’-Cy3/Cy5-oligo(dT)_15_-dsDNA construct with 3’ polarity as a function of protein concentration ([gp32] = 0.0, 0.1, 0.5 and 1.0 *μ*M). We also examine the sensitivity of the assembly pathway to the length and polarity of the 3’,5’-oligo(dT)_14,15_ lattice at fixed protein concentration ([gp32] = 0.5 *μ*M). We summarize the results of these studies in terms of the measured probability distribution functions (PDFs), the two-point time correlation functions (TCFs) and the three-point TCFs in Fig. 2. Theoretical considerations involved in simulating the PDFs and TCFs using the kinetic network model, as discussed further below, are summarized in the SI section. Equivalent studies of the assembly of the first two gp32 monomers onto DNA constructs with polarities 3’ and 5’ and with lattice length *n* = 14 nts are also described in the SI section of this paper, since the same experimental approaches were used in all these studies, though the results differed. The effects of ssDNA composition (here oligo-dT sequences), lattice polarity (5’ or 3’) and lattice length (*n* = 14 or 15 nts) are then compared in the Discussion section of the main text (see below).

**Figure 2.**
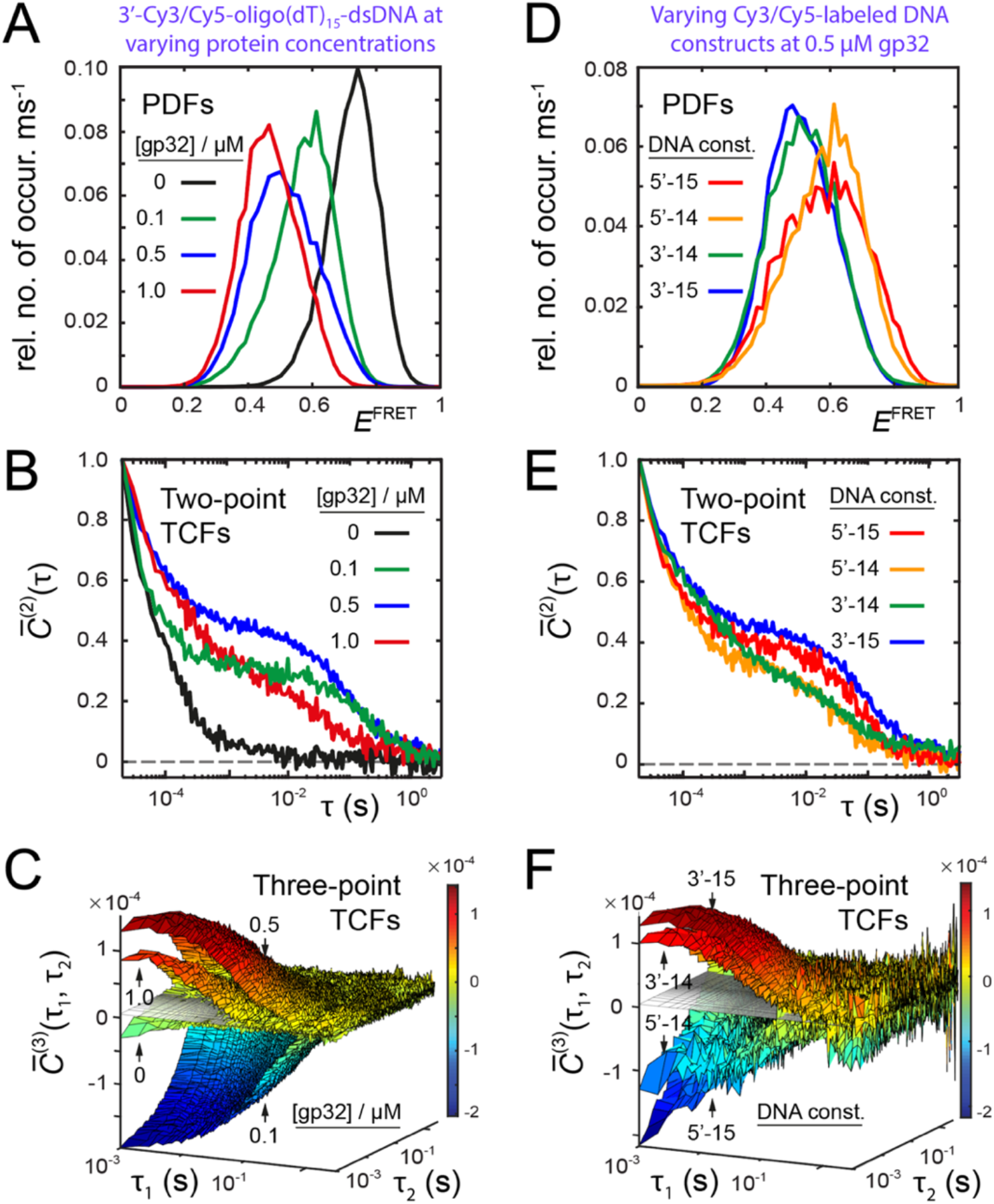
Results of smFRET studies for (***A*** - ***C***) the 3’-Cy3/Cy5-oligo(dT)_15_-dsDNA construct as a function of ssb protein concentration ([gp32] = 0, 0.1, 0.5 and 1.0 *μ*M), and (***D*** - ***F***) the four Cy3/Cy5-oligo(dT)_*n*_-dsDNA constructs at [gp32] = 0.5 *μ*M. The DNA constructs are listed in Table 1 and vary in oligo(dT)_*n*_ lattice length (*n* = 14 or 15 nts) and polarity (3’ or 5’). The functions shown are: (***A, D***) the probability distribution functions (PDFs), which were constructed using a signal integration period, *T*_*w*_ = 5 ms; (***B, E***) the two-point time-correlation functions (TCFs) that were constructed using *T*_*w*_ = 10 *μ*s; and (***C, F***) the three-point TCFs that were constructed using *T*_*w*_ = 1 ms. The horizontal gray surfaces shown in panels C and F serve as positional reference indicators.

In Figs. 2*A* – 2*C*, we compare our results for the 3’-Cy3/Cy5-oligo(dT)_15_-dsDNA construct at varying DNA construct and varying protein concentrations: [gp32] = 0.0, 0.1, 0.5 and 1.0 *μ*M. We first consider the concentration-dependence of the PDFs, which are normalized histograms of the *E*^*FRET*^ values (see Fig. 2*A*). In the absence of protein, the PDF (shown in black) is asymmetric with its peak value at 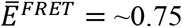 and a shoulder extending to lower *E*^*FRET*^ values, indicating the presence of a non-uniform distribution of conformational species. In prior studies we obtained similar results for these same DNA constructs (32), which revealed that in the absence of protein the 3’-oligo(dT)_15_ lattice interconverts on sub-millisecond time scales between distinct conformational macrostates that differ between lattice chain extensions and the extent of unstacking between adjacent nts. At monomer protein concentrations ([gp32] = 0.1, 0.5 and 1.0 *μ*M), the peaks of the PDFs (green, blue and red, respectively) shift systematically to lower *E*^*FRET*^ values (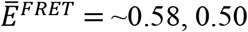 and 0.47), indicating that gp32 proteins are binding to and stabilizing extended conformational macrostates of the 3’-Cy3/Cy5-oligo(dT)_15_-dsDNA construct. As the protein concentration is increased, the relative populations of conformational macrostates with lower *E*^*FRET*^ values (and correspondingly longer chain extensions) are also increased.

The dynamics of the conformational macrostates are characterized by the two-point TCFs (see Fig. 2*B*). In the absence of protein, the two-point TCF decays rapidly to zero on sub-millisecond time scales (black curve) (32). However, at elevated protein concentrations the dynamics slow dramatically; the additional decay components span a broad range of time scales from hundreds-of-microseconds to hundreds-of-milliseconds. At the lowest protein concentration (0.1 *μ*M, green) the fastest decay component (≤ 100 *μ*s) closely resembles that of the DNA construct in the absence of protein. However, on longer time scales the TCF exhibits a ‘plateau’ region from 1 – 100 ms followed by a ‘slow’ decay at ∼ 300 ms. At higher protein concentrations (0.5 *μ*M, blue; 1.0 *μ*M, red), the fastest decay components slow significantly, and at the highest protein concentration (1.0 *μ*M) the slowest decay component speeds up to ∼10 ms.

Additional information about the details of the transition pathways is contained in the three-point TCFs. As shown in Fig. 2*C*, the amplitudes and signs of the three-point TCFs at short times (*T*_*w*_ = ∼1 ms) are sensitive to the concentration of protein. This is due to concentration-dependent changes in the populations of protein-bound versus unbound macrostates (as indicated by the PDFs, see Fig. 2*A*) and differences in the prevalent transition pathways at short times. For the 3’-Cy3/Cy5-oligo(dT)_15_-dsDNA construct in the absence of protein, the three-point TCF exhibits relatively weak negative amplitude. However, at low protein concentration (0.1 *μ*M) the three-point TCF exhibits strong negative amplitude. When the protein concentration is increased (to 0.5 *μ*M), the amplitude of the three-point TCF is strongly positive, and at even higher concentration (1.0 *μ*M) the sign remains positive, but its amplitude is somewhat diminished. We further discuss the protein concentration-dependence of the PDFs, two-point TCFs and three-point TCFs in the following sections in which we present our analyses of these results.

We next compare our smFRET data for the four [3’,5’-Cy3/Cy5-oligo(dT)_14,15_-dsDNA] constructs listed in Table 1 at [gp32] = 0.5 *μ*M. In Fig. 2*D*, we present the PDFs for the four constructs, which we designate as 3’-15, 3’-14, 5’-15 and 5’-14, indicating the polarities (3’ and 5’) and lengths (*n* = 14 and 15 nts) of the oligo(dT)_*n*_ in relation to the positions of the ss-dsDNA junctions. We note that the PDF for the 3’-15 construct (shown in blue) is the same as that in Fig. 2*A*, which is approximately symmetric about its peak value, 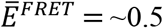. For the 3’-14 construct, the PDF (dashed green) is very similar to that of the 3’-15 construct with approximately the same peak value. In contrast, the PDFs for the 5’-15 and 5’-14 constructs (red and orange, respectively) are more disperse than their 3’ counterparts and their peaks are shifted to significantly higher peak positions, 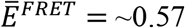. Thus, the equilibrium populations of protein-bound states (with longer chain extensions) are greater for the 3’-15 and 3’-14 constructs than for the 5’-15 and 5’-14 constructs.

State-to-state interconversion dynamics are characterized by the two-point TCFs shown in Fig. 2*E*. All four constructs exhibit distinct kinetic properties. The 3’-15 construct (blue) exhibits the slowest decay on all time scales (including a ‘quasi-plateau’ region from 1 – 100 ms), while the 5’-14 construct (orange) exhibits the fastest overall decay. For the 3’-14 construct (green), the fastest decay component (≤ 100 *μ*s) is like that of the 3’-15 construct, however the 3’-14 construct does not exhibit a ‘quasi-plateau.’ Rather, the 3’-14 construct exhibits faster decay components on the intermediate time scales of ∼1 – 100 ms. For the 5’-15 construct (red), the sub-millisecond decay component is even faster than those of the 3’-15 and 3’-14 constructs. However, the 5’-15 construct does exhibit a ‘plateau’ region at intermediate time scales like that of the 3’-15 construct. As noted above, the 5’-14 construct exhibits the fastest overall dynamics. At sub-millisecond time scales, the dynamics of the 5’-14 construct resemble those of the 5’-15 construct, but at intermediate time scales the 5’-14 construct resembles the 3’-14 construct. In summary, 3’,5’-oligo(dT)_14,15_-dsDNA constructs with the same polarity exhibit similar sub-millisecond dynamics (5’ being faster and 3’ being slower) while constructs with the same lengths exhibit similar intermediate time scale dynamics (*n* = 14 nts being faster and *n* = 15 nts being slower).

Additional information about the dynamics of these systems is contained in the three-point TCFs, shown in Fig. 2*F*. These data show that the short-time amplitudes and signs of the three-point TCFs are sensitive to the polarities and lattice lengths of the four constructs. For the 3’-15 and 3’-14 constructs, the amplitudes are strongly positive. However, the 3’-14 construct exhibits slightly weaker amplitude than the 3’-15 construct. The situation is opposite for the 5’-15 and 5’-14 constructs, which both exhibit strongly negative amplitudes with the 5’-14 construct being slightly weaker than the 5’-15 construct. In the following sections, we further discuss the length and polarity dependence of the PDFs, two-point TCFs, and three-point TCFs in the context of our theoretical analyses and interpretations of these functions.

We analyzed our data using the kinetic network model that is outlined in the SI section and described in our previous work (24, 32-34). For all the conditions that we studied, we determined that our data are best modeled using a five-state network scheme. This is supported by the fact that all the two-point TCFs are well described as a sum of four single exponential decay components (neglecting the slow mechanical noise component) with time constants separated from one another by an order of magnitude in each case (see Fig. S1 and Table S1 of the SI).

We summarize our results for the 3’-oligo(dT)_15_-dsDNA construct as a function of protein concentration in Fig. 3. The schemes corresponding to [gp32] = 0.1, 0.5 and 1.0 *μ*M, respectively (Figs. 3*A* – 3*C*) are based on our optimization results shown in Fig. S2 of the SI. These schemes resulted from a thorough exploration of the parameter space in which we tested hundreds of different network configurations in parallel using a high-performance computer cluster. The five macrostates are labeled, *S*_1_ – *S*_5_, in order of decreasing *E*^*FRET*^ values, as shown (optimized parameters listed in Table S2 of the SI). Also shown are the optimized values of the equilibrium populations, 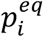, the signal fluctuations, 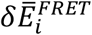 (Table S3 of the SI), and the forward and backward time constants, 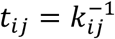 (Table S4 of the SI). From the optimized equilibrium and kinetic parameters, we determined the free energy minima (Table S5 of the SI) and activation energies (Table S6 of the SI) using the Boltzmann and Arrhenius equations, respectively. We have thus approximated the FES diagrams shown in Figs. 3*D* – 3*F*.

**Figure 3.**
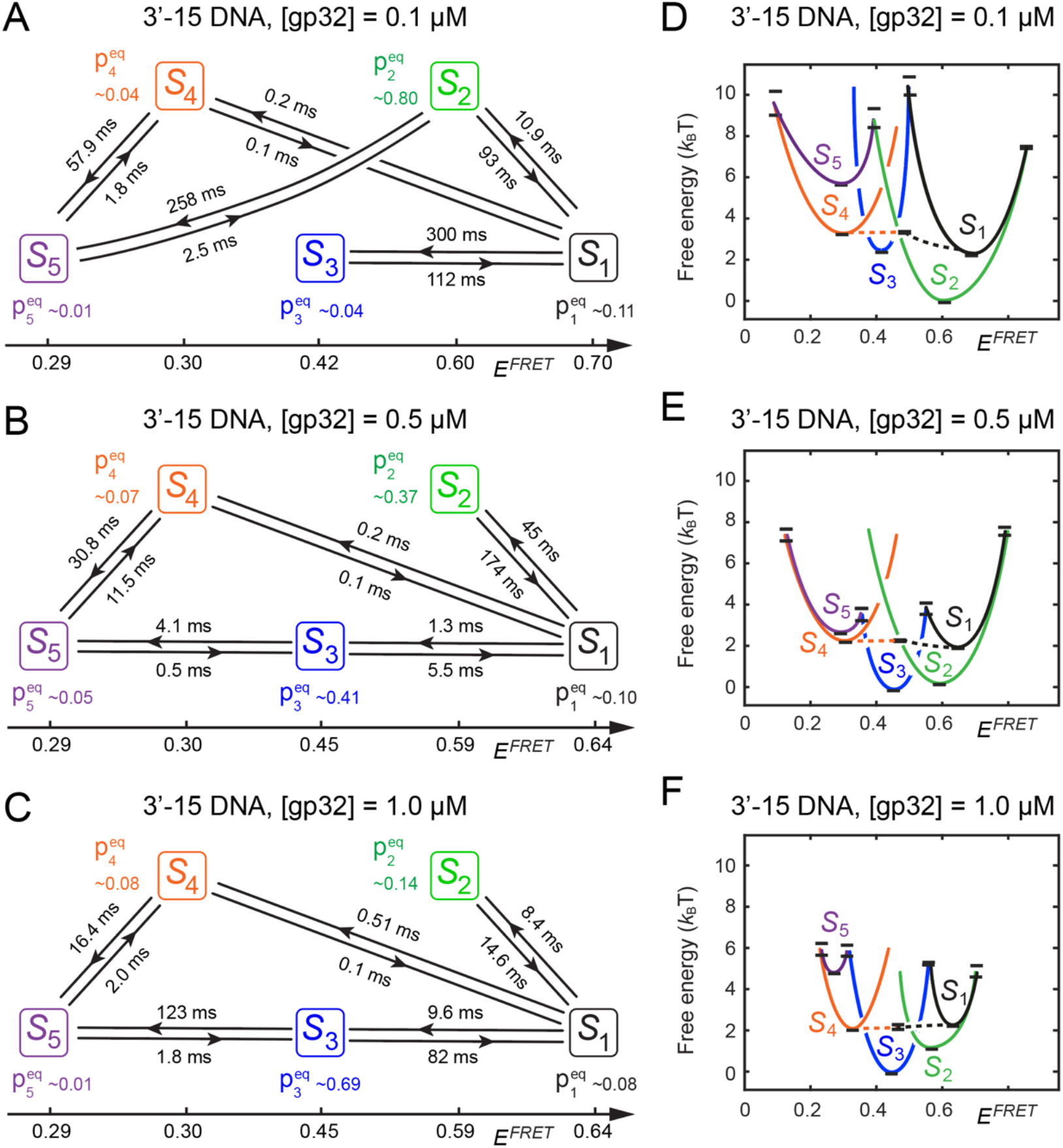
Kinetic network schemes and FESs for the 3’-oligo(dT)_15_-dsDNA construct in the presence of (***A, D***) [gp32] = 0.1 *μ*M, (***B, E***) 0.5 *μ*M and (***C, F***) 1.0 *μ*M in aqueous buffer with 100 mM NaCl and 6 mM MgCl_2_. Panels (A) - (C) show the network schemes and optimized values for the equilibrium populations, 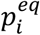, time constants, 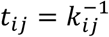, and mean FRET efficiencies, 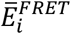, which were determined from the analysis of microsecond-resolved smFRET data. Structural assignments for the five macrostates are: *S*_1_ (the ‘free’ DNA construct); *S*_2_ (a gp32 monomer-DNA-bound conformation at a medial position within the 3’-oligo(dT)_15_ lattice at positions 3 – 9, 4 – 10, …, 7 – 13); *S*_3_ (a gp32 monomer-DNA-bound conformation at the ss-dsDNA junction at positions 1 – 7 or 2 – 8 near the ss-dsDNA junction); *S*_4_ (a gp32 monomer-DNA-bound conformation at the 3’ distal end at positions 8 – 14 or 9 – 15); and *S*_5_ (one of two possible gp32 dimer conformations (at positions 1 – 14 or 2 – 15). Panels D – F show the corresponding FESs inferred from the free energy minima and activation barriers, which were calculated from the optimized equilibrium and kinetic parameters using the Boltzmann and Arrhenius equations, respectively (details provided in the SI section).

In Table S7 of the SI, we list (in order of descending magnitude) the twenty largest positive and negative pathway terms that contribute to the two-point TCF [*i*.*e*., the 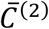 function] at *τ* = 10 μs for each of the DNA constructs and sample conditions studied in this work. Each term is assigned to a specific two-point pathway (*i* → *j*) and factored into its two-point product of signal fluctuations, 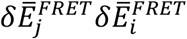, and its assigned statistical weight, 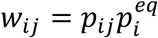 [see Eq. (6)]. We thus find that the 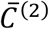 function is most heavily weighted by positive terms like 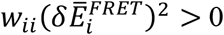, in which the initially measured macrostate, *S*_*i*_, is measured again within the relatively short period *τ* = 10 μs. This is because the survival probabilities, *p*_*ii*_, are generally large for small *τ*, as shown in the fourth row of Fig. S2 of the SI (solid curves). In Table S8 of the SI, we list for each of the samples the twenty largest positive and negative terms (of 125 total) that contribute to the three-point TCF [the 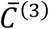 function] at *τ*_1_ = *τ*_2_ = 1 ms. In this case, each term is assigned to a specific three-point pathway (*i* → *j* → *k*) and factored into its three-point fluctuation product, 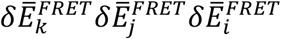, and statistical weight, 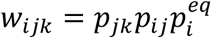 [see Eq. (7)]. These three-point pathway terms are useful to interpret the signs and relative magnitudes of the experimentally-derived 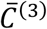 functions shown in the third rows of Fig. S2 and Fig. S3 of the SI. For example, the 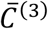 function at [gp32] = 0.1 *μ*M is dominated by the negative term 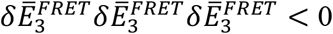. At this low protein concentration, pathway terms involving multiple factors of 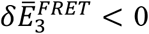 are relatively large due to the large survival probability, *p*_33_ (see Fig. S2*A* of the SI, bottom row). This contrasts with the situation at higher protein concentrations ([gp32] = 0.5 *μ*M and 1.0 *μ*M, Figs. S2*B* and S2*C* of the SI, respectively) for which the dominant terms are positive: *e*.*g*., 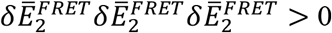.

We made the following assumptions in assigning macrostates *S*_1_ – *S*_5_ to the relevant end-states and intermediates involved in the assembly mechanism. In all cases, we assumed that macrostate *S*_1_ (with the highest *E*^*FRET*^ value) corresponds to a ‘compact’ chain conformation in which no gp32 protein is bound to the 3’-oligo(dT)_15_-dsDNA construct. Conversely, macrostate *S*_5_ (with lowest *E*^*FRET*^ value) corresponds to a ‘highly extended’ chain conformation in which two gp32 proteins are bound cooperatively as a dimer to the DNA construct. Macrostates *S*_2_, *S*_3_ and *S*_4_ (with *E*^*FRET*^ values intermediate to those of *S*_1_ and *S*_5_) each corresponds to a unique gp32 monomer-DNA-bound conformation that can be generated from macrostate *S*_1_. Of the three monomer-DNA-bound conformations, two serve as ‘productive’ intermediates that proceed directly to generate the gp32 dimer-DNA-bound macrostate, *S*_5_, while the third monomer-DNA-bound intermediate represents a ‘dead-end’ species that is not on the pathway to form *S*_5_.

The network schemes shown in Figs. 3*A* – 3*C* indicate that for each of the protein concentrations that we studied, a single monomer-DNA-bound intermediate of significant population dominates the assembly mechanism. These are: *S*_1_ ↔ *S*_2_ ↔ *S*_5_ for [gp32] = 0.1 *μ*M and *S*_1_ ↔ *S*_3_ ↔ *S*_5_ for [gp32] = 0.5 and 1.0 *μ*M. These predominant pathways can also be discerned by examining the FESs shown in Figs. 3*D* – 3*F*, which indicate that the intermediate *S*_2_ is most stable for [gp32] = 0.1 *μ*M and *S*_3_ is most stable for [gp32] = 0.5 and 1.0 *μ*M. We note that for all the protein concentrations the intermediate *S*_4_, which rapidly interconverts with *S*_1_ on sub-millisecond time scales, is sparsely populated 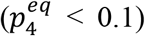. Thus, *S*_1_ ↔ *S*_4_ ↔ *S*_5_ represents a relatively minor pathway in the overall mechanism. Since the distal end of the 3’-oligo(dT)_15_ lattice can fluctuate rapidly to present accessible binding sites for gp32 monomers in solution, we assign the *S*_4_ macrostate to the monomer-DNA-bound complex at the distal end of the 3’-oligo(dT)_15_ lattice.

We next consider our assignments for the gp32 monomer-DNA-bound macrostates *S*_2_ and *S*_3_. At the lowest tested protein concentration ([gp32] = 0.1 *μ*M, Fig. 3*A*), macrostate *S*_2_ is the majority component species 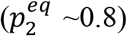 that forms relatively quickly (∼11 ms). Moreover, *S*_2_ is kinetically stable with mean lifetime (*k*_21_ + *k*_25_)^−1^ = ∼68 ms. Conversely, at this low protein concentration the *S*_3_ macrostate forms relatively slowly (∼300 ms) and is sparsely populated 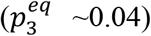– *i*.*e*., *S*_3_ is unstable relative to *S*_2_ (see Fig. 3*D*). Because there are multiple binding sites available within the medial regions of the 3’-oligo(dT)_15_ lattice, we assign the *S*_2_ macrostate to a gp32-monomer-DNA-bound complex at any of the medial positions. During the ∼68 ms lifetime of the *S*_2_ macrostate, the gp32 monomer may slide along the 3’-oligo(dT)_15_ lattice to a position close to the distal 3’ end or the ss-dsDNA junction to permit a second gp32 monomer to bind and form the gp32 dimer-DNA-bound complex, *S*_5_.

As the protein concentration is increased to [gp32] = 0.5 *μ*M and then to 1.0 *μ*M, the *S*_3_ macrostate becomes progressively more stable while the *S*_2_ macrostate becomes progressively less stable (see Figs. 3*E* and 3*F*). Moreover, the transition barriers for the *S*_1_ ↔ *S*_3_ ↔ *S*_5_ pathway are reduced relative to those for the *S*_1_ ↔ *S*_2_ ↔ *S*_5_ pathway at [gp32] = 0.1 *μ*M. At these elevated protein concentrations, the gp32 monomer is more likely to encounter the 3’-Cy3/Cy5-oligo(dT)_15_-dsDNA construct at multiple sites and with multiple protein conformations (see Fig. 1*A*), leading to the rapid formation of the *S*_3_ intermediate. We thus assign the *S*_3_ macrostate to the gp32 monomer-DNA-bound complex at the ss-dsDNA junction, which once formed converts rapidly to the cooperatively-bound dimer-DNA complex. We note that the transition barriers for the assembly pathway *S*_1_ ↔ *S*_3_ ↔ *S*_5_ are significantly lower at [gp32] = 0.5 *μ*M in comparison to [gp32] = 1.0 *μ*M. The increased barrier heights at the higher protein concentration may reflect the presence of multiple gp32 monomers competing for the available DNA binding sites, and thus occluding the lattice with gp32 monomers bound at central lattice positions, which decreases the availability of lattice sites at the ss-dsDNA junction (1 – 7 or 2 – 8) from which cooperatively bound gp32 dimers can form.

We next examine the dependence of the gp32 assembly mechanism on the four DNA constructs listed in Table 1, which vary in oligo(dT)_*n*_ lattice length (*n* = 14, 15 nts) and polarity (3’ versus 5’). For all four DNA constructs at [gp32] = 0.5 *μ*M, we find that the mechanism is best described using the same five-state kinetic scheme as the one discussed above for the 3’-oligo(dT)_15_-dsDNA (3’-15) construct. In Fig. 4 we compare our results for the 3’-15 construct to those of the 5’-15 construct, and in Fig. S4 of the SI for the 3’-14 construct to those of the 5’-14 construct. A detailed comparison between our simulated and experimental functions is shown in Fig. S3 of the SI. As mentioned previously, the three-point TCFs exhibit positive amplitudes for the 3’-15 and 3’-14 DNA constructs and negative amplitudes for the 5’-15 and 5’-14 DNA constructs. The three-point pathway terms listed in Table S8 of the SI show that the dominant contributions to the 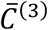 functions at *τ*_1_ = *τ*_2_ = 1 ms are 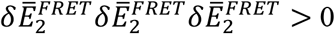 for the 3’-15 construct, 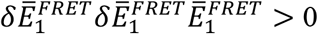 for the 3’-14 construct, and 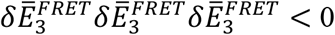 and 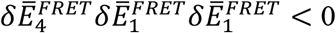 for the 5’-15 and 5’-14 constructs, which is in keeping with the signs of the short time amplitudes of these functions.

**Figure 4.**
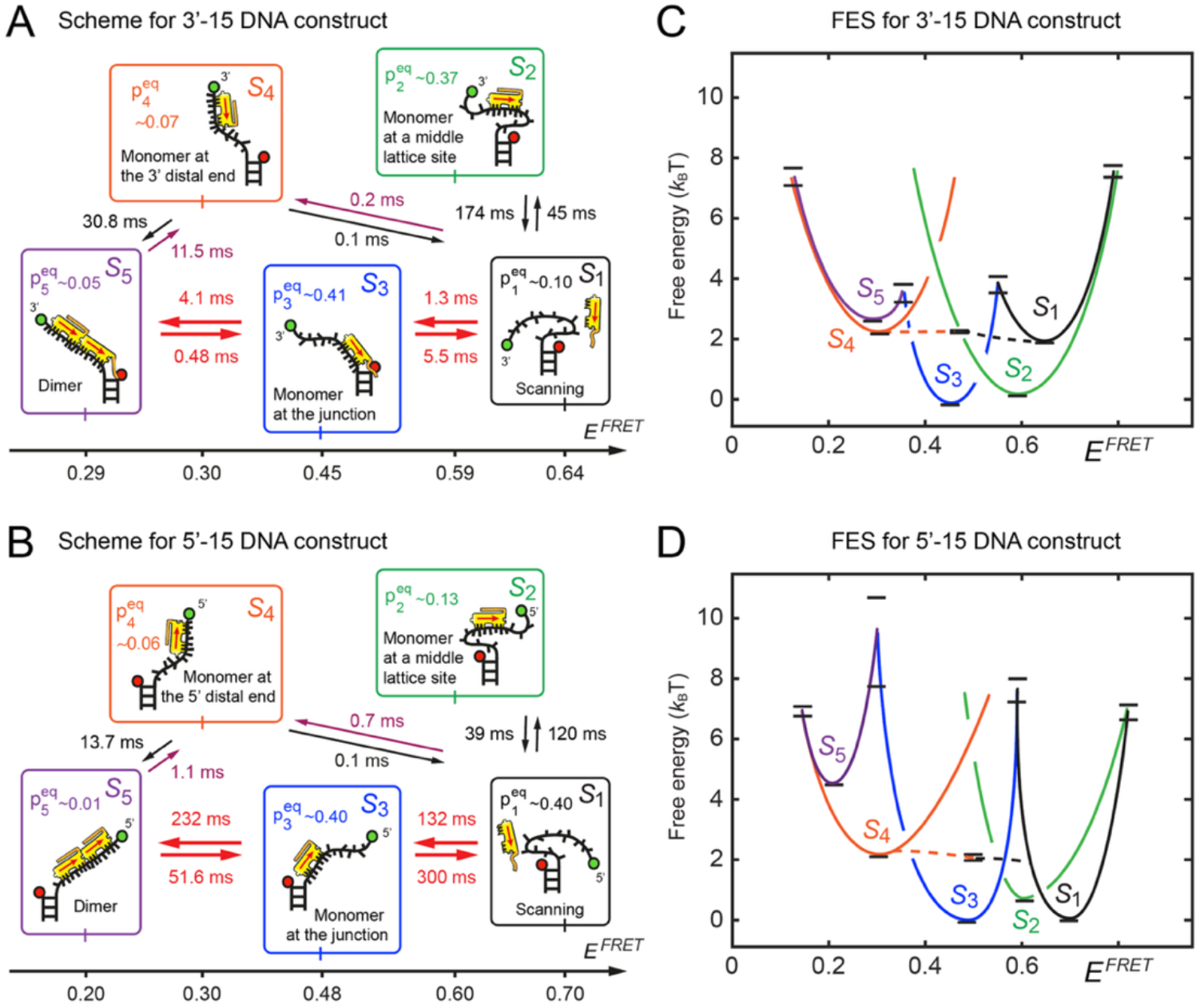
Kinetic network schemes and FESs for (***A, C***) the 3’-oligo(dT)_15_-dsDNA construct (3’-15) and (***B, D***) the 5’-oligo(dT)_15_-dsDNA construct (5’-15) in the presence of [gp32] = 0.5 *μ*M in aqueous buffer containing 100 mM NaCl and 6 mM MgCl_2_. Optimized kinetic and equilibrium parameters are listed, as in Fig. 3. The structural assignments for macrostates *S*_1_ - *S*_5_ are depicted schematically indicating the orientation of the CTD relative to the ss-dsDNA junction and the polarity of the oligo(dT)_15_ overhanging tail.

We find that the equilibrium and kinetic parameters for the 3’-15 DNA construct are very different from those for the 5’-15 construct. As discussed in the previous section, the primary assembly pathway for the 3’-15 construct is *S*_1_ ↔ *S*_3_ ↔ *S*_5_ whose transition barriers are relatively small (∼4 *k*_B_*T*, see Table S6 of the SI and Figs. 4*A* and 4*C*). Furthermore, for the 3’-15 construct the unbound species *S*_1_ is weakly populated 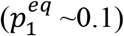 in comparison to the monomer-DNA-bound states 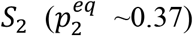 and 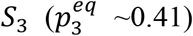. We interpret these findings using the structural assignments presented in Fig. 4*A*, in which the *S*_1_ macrostate is depicted with the CTD of the gp32 monomer extended towards duplex region of the ss-dsDNA junction in a ‘scanning’ conformation. Here, the favorable interaction between the CTD and the monovalent cation condensation layer that stabilizes the duplex region serves to orient the gp32 monomer relative to the 3’-ss-dsDNA junction (as depicted in Fig. 1*B*), thus facilitating the initial docking and subsequent formation of the *S*_3_ macrostate. In contrast, for the 5’-15 construct the transition barriers of the *S*_1_ ↔ *S*_3_ ↔ *S*_5_ pathway are relatively high (∼8 *k*_B_*T*) and the unbound macrostate *S*_1_ is a majority species 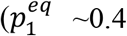, Figs. 4*B* and 4*D*). In keeping with the structural assignments that we adopted above, Fig. 4*B* shows the *S*_1_ macrostate with the CTD of the gp32 monomer extended towards the ss-dsDNA junction. However, to form the monomer-DNA-bound macrostate *S*_3_ the orientation of the protein must be reversed. In addition, we note that the ‘unproductive’ intermediate *S*_2_ is much less populated for the 5’-15 construct 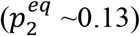 than for the 3’-15 construct 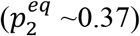, suggesting that the role of the CTD to orient the protein relative to the ss-dsDNA junction may also influence the transition barrier and stability of the *S*_2_ macrostate.

We find the behavior of the *n* = 14 nts constructs to be very similar to that of the *n* = 15 nts constructs (see Fig. S4 of the SI). The primary assembly pathway is *S*_1_ ↔ *S*_3_ ↔ *S*_5_, and the transition barriers are significantly lower for the 3’-14 versus the 5’-14 construct. The effect of decreasing lattice length for the 3’ polarity constructs by a single nucleotide is to increase the transition barrier for the *S*_1_ ↔ *S*_3_ step (from ∼4 *k*_B_*T* to ∼6 *k*_B_*T*) while leaving the barrier for the *S*_1_ ↔ *S*_2_ step largely unaffected (Table S6 of the SI). Moreover, the ‘unproductive’ *S*_2_ macrostate becomes destabilized relative to the ‘productive’ *S*_3_ macrostate (from ∼0.3 *k*_B_*T* to ∼0.6 *k*_B_*T*, see Table S5 of the SI). These effects may be due to the oligo(dT)_14_ lattice adopting a limited number of conformations that are suitable for the gp32 monomer (correctly oriented at the ss-dsDNA junction) to dock and bind at the single binding site (1 – 7) near the ss-dsDNA junction to form the *S*_3_ macrostate.

It is interesting to note that the effects of lattice length on the 5’ polarity constructs are opposite to those of the 3’ polarity constructs. For the 5’-14 construct, the transition barriers for the *S*_1_ ↔ *S*_3_ and *S*_1_ ↔ *S*_2_ steps are smaller than those of the 5’-15 construct (∼ 5 *k*_B_*T* versus ∼7 *k*_B_*T*). Furthermore, the ‘unproductive’ monomer-DNA-bound species *S*_2_ becomes more stable than the ‘productive’ monomer-DNA-bound species *S*_3_ (compare Figs. S4*C* and S4*D* of the SI). These effects may be due to conformations adopted by the oligo(dT)_14_ lattices that favor the docking and binding of the gp32 monomer at the multiple middle binding sites of the oligo(dT)_14_ lattice, rather than at the single gp32 binding site (1 – 7) near the ss-dsDNA junction.

In the discussion that follows we attempt to put the above findings into a more general and biologically relevant context.

## IV. Discussion

In this work, we have applied microsecond-resolved smFRET to study the initial nucleation steps of the T4 bacteriophage gp32 (ssb) protein onto model oligo(dT)_*n*_-dsDNA constructs of varying lattice length (*n* = 14 nts versus *n* = 15 nts) and polarity (3’ versus 5’). We analyzed our data using a previously developed method that compares probability distribution functions (PDFs) and time correlation functions (TCFs) to determine the kinetic and equilibrium parameters of a five-state kinetic network scheme (24, 32-34).

Our results lead us to the following conclusions: *i*) the initial nucleation and assembly steps of the gp32 protein are highly sensitive to the polarity of the oligo(dT)_*n*_ lattice relative to the ss-dsDNA junction; *ii*) for the 3’-15 construct, the gp32 monomer-DNA-bound intermediate forms at the ss-dsDNA junction (macrostate *S*_3_) with a significantly lower activation barrier (*t*_13_ ∼1.3 ms) and greater stability (Δ*G*_13_ ≈ −1.4 k_B_*T*) than displayed by the 5’-15 construct (*t*_13_ ∼132 ms, Δ*G*_13_ ≈ 0); *iii*) the rates of these initial steps (though not their relative stabilities) are weakly dependent on lattice length, with the 3’-14 construct being slightly slower to form the 3’-15 construct (*t*_13_ ∼7.2 ms versus ∼1.3 ms), and the 5’-14 construct forming slightly faster than the 5’-15 construct (*t*_13_ ∼48.1 ms versus ∼132 ms).

The polarity-dependence of the gp32 binding and assembly mechanism has been addressed in previous work (13, 14, 16, 17, 20, 31), although most previous studies have focused on the gp32 binding affinities at equilibrium rather than on the kinetics of the binding process. We note that the earlier work from our laboratory did not enjoy the microsecond time resolution that we have now achieved with our present experimental setup. Our conclusions are largely consistent with the results of prior studies from our laboratories that utilized spectroscopic measurements performed under ‘physiological’ salt concentration conditions (16, 17, 20, 31), while they are not consistent with other studies in which gel electrophoresis measurements were utilized to estimate binding affinities (13, 14). It is likely that these discrepancies are due to the difficulties of establishing defined salt concentrations in gel electrophoresis experiments. Indeed, the molecular mechanisms that we propose to interpret our results are strongly dependent on the conformations of the gp32-fork junction complexes and the interactions between the CTD and the DNA construct, both of which are highly sensitive to salt concentration.

In addition to the salt concentration-gel electrophoresis problem alluded to above, it is possible that the discrepancies between our results and the related measurements of Jones *et al*. (14) and Jordan *et al*. (13) could also (in part) reflect differences in the DNA topologies. These workers carried out their experiments on model DNA constructs containing ‘flap junctions,’ in which the ssDNA leading or lagging strand arms of a model ss-dsDNA fork construct were alternately hybridized using complementary strands. However, and in any case, the salt anomalies associated with gel electrophoresis experiments of this type make our data and theirs difficult to compare.

The nucleation mechanism for initial gp32 cluster formation that we have proposed involves a ‘pre-binding orientation’ step that favors binding of gp32 monomers close to ss-dsDNA junctions of 3’ constructs and disfavors binding close to the junctions of 5’ constructs. We posit that the negatively charged CTD in the extended conformation of the gp32 monomer is attracted to the relatively dense condensation layer of monovalent sodium ions near the surface of the duplex region of the ss-dsDNA junction (see Fig. 1*A*). Thus, for the 3’ constructs, the CTD of the gp32 monomer of macrostate *S*_3_ points towards the ss-dsDNA junction, while for the 5’ constructs, the CTD of macrostate *S*_3_ points away from the junction. An alternative scenario is that the CTD preferentially interacts with the distal 3’ or 5’ ends of oligo(dT)_14,15_ lattices. However, our proposed nucleation mechanism, which is based on the electrostatic interactions between the CTD and the sodium ion concentration gradient at ss-dsDNA junctions, seems more plausible. Future studies using gp32 mutants in which the CTD is deleted or otherwise altered could reaffirm or disprove our proposed model.

The biological significance of our model studies of the initial nucleation steps of ssDNA-gp32 filament *assembly* is extended by comparing our findings with those of Cashen *et al*. (9, 10) that deals with the other end of the process – *i*.*e*., the *disassembly* of the fully formed ssDNA-gp32 filaments, as illustrated in Fig. 5. Here we schematize our proposed mechanism for gp32-ssDNA filament assembly at the primer-template ssRNA-ssDNA junction and the mechanism put forward by Cashen *et al*. for sequential disassembly of cooperatively bound gp32 monomers from the previously-formed gp32-ssDNA filament due to *lagging strand* DNA synthesis. As the primer-template DNA junction emerges from the replication fork, a gp32 monomer carrying an extended CTD orients itself (and its CTD) relative to the junction (Fig. 5*A*) before forming a non-cooperatively bound gp32-ssDNA complex (Fig. 5*B*). Upon further elongation and exposure of additional lattice sites, gp32 monomers bind successively (Fig. 5*C*) to form a cooperatively bound (gp32)_*n*_-ssDNA filament. Of course, it is possible that the initial gp32 monomer orientation and binding steps at the primer-template DNA junction may involve additional interactions between gp32 and the gp45 sliding clamp immediately after the clamp is loaded by the clamp-loader complex (5). This possibility will be investigated in future studies. We note that the time scale of filament assembly that we have determined at the lagging strand p/t-DNA junction (*i*.*e*., a few milliseconds per gp32 monomer, see Fig. 4*A*) matches the actual rate of DNA synthesis (∼1,000 nts s^-1^) in T4 bacteriophage infected *E. coli* cells. We emphasize that we were able to accurately measure these millisecond processes by employing our microsecond-resolved smFRET method and analysis (24, 32, 34).

**Figure 5.**
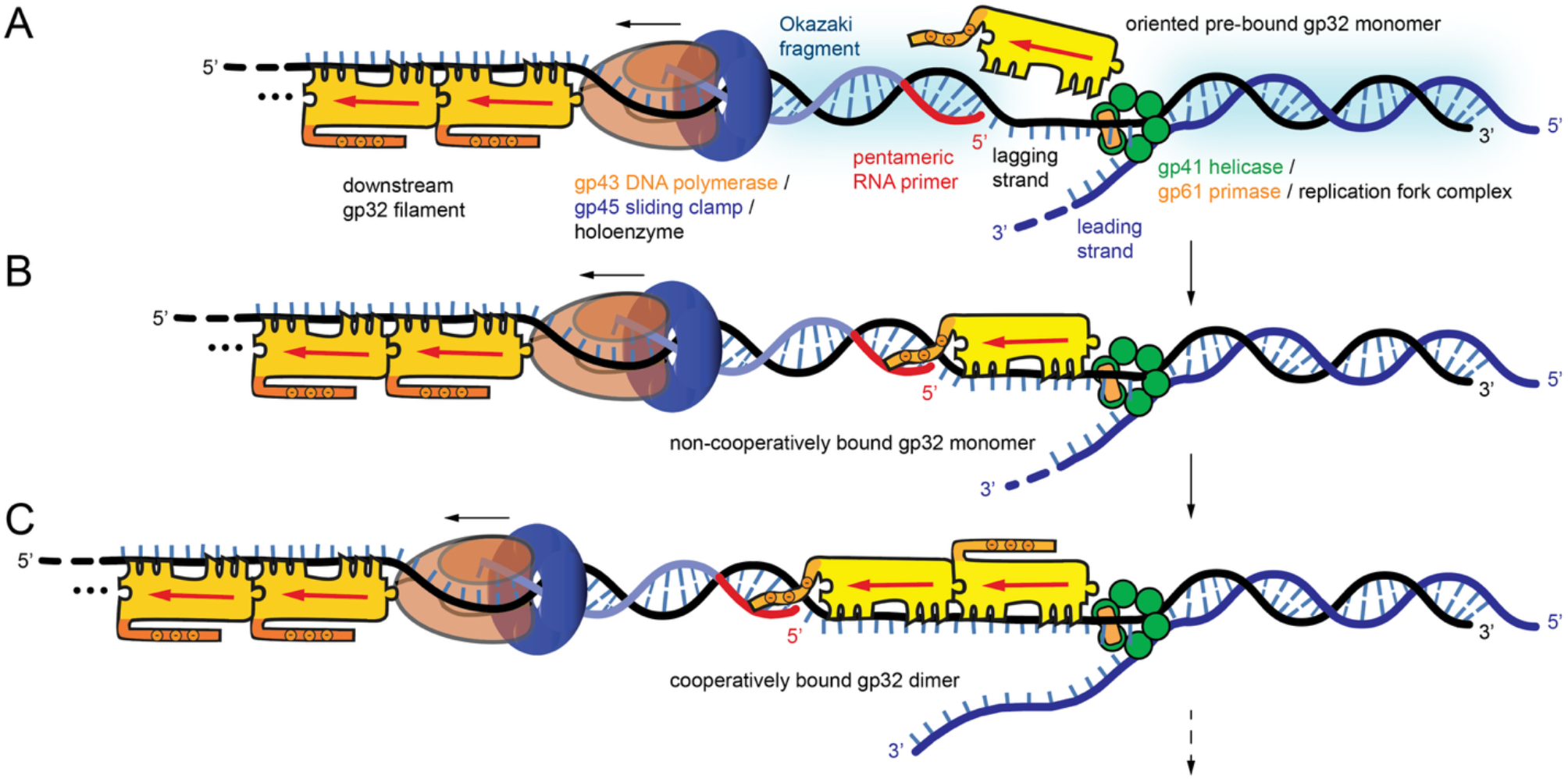
Proposed mechanism for the nucleation and cooperative binding steps of gp32 ssb proteins (shown in yellow) during the elongation cycle of the T4 bacteriophage DNA replication system. This is compared to the disassembly mechanism proposed by Cashen *et al*. (9, 10). (***A***) At the replication fork the gp41 replication helicase (green) unwinds the genomic DNA, thus exposing leading (black) and lagging (blue) DNA template strands at a rate of ∼1,000 nts per second. A pentameric RNA primer sequence (red) is synthesized on the lagging strand by the gp61 primase, which is the starting point for the synthesis by the DNA polymerase holoenzyme of a newly formed Okazaki fragment (purple). As the lagging strand is fed out from the central channel of the helicase, a gp32 monomer with extended CTD orients relative to the primer-template (ssRNA-ssDNA) junction in a ‘pre-binding’ conformation. Light blue shaded regions indicate the cation condensation layer near the surfaces of the duplex regions. (***B***) When this process has fully exposed a 7 nt binding site on the lagging strand template, the gp32 protein monomer binds to the template in a non-cooperative modality. (***C***) Upon further elongation, gp32 proteins bind successively in a cooperative process. The final two monomers of the filament from the previous cycle (darker yellow) are shown downstream of the lagging strand polymerase. Filament disassembly occurs by a compression process coupled to DNA synthesis in which cooperative interactions between adjacent gp32 monomers are disrupted, thus destabilizing the downstream filament (9, 10).

The role of the CTD in our proposed assembly mechanism is complementary to the filament disassembly mechanism recently described by Cashen *et al*. (9, 10) These workers performed force-extension (optical tweezer) experiments on long gp32 filaments (∼ 8,000 nts), which showed that the presence of the full CTD is required to disrupt cooperative interactions between adjacent gp32 monomers. As the Okazaki fragment is synthesized by the DNA polymerase holoenzyme (see Fig. 5), the downstream gp32 filament is compressed and cooperative protein-protein interactions are weakened, ultimately leading to gp32 monomer dissociation and filament disassembly. Our current results serve to complete this picture by suggesting a versatile role for the CTD in mediating *both* the assembly and disassembly processes of the gp32-ssDNA filaments that are central to overall function of the replisome system. We note that a recent paper by Xu *et al*. (36) suggests that the DNA polymerase of the T7 bacteriophage DNA replication system also actively and sequentially displaces the T7 single-stranded DNA binding (ssb) proteins.

In addition to their significance for DNA replication, our findings provide unique insights into the initial nucleation and assembly steps of gp32 proteins at various types of ss-dsDNA junctions. The possible role played by the CTD of gp32 in the polarity-dependent recognition of ss-dsDNA junctions is relevant to other recognition processes in molecular biology. For example, regulatory proteins often contain intrinsically disordered regions (IDRs), which are thought to participate in the search for, and identification of, promotor and operator sites (37). Moreover, such search and recognition events are likely important in controlling the dynamics of chromatin movement in higher organisms, functioning to expose or conceal relevant transcription factor binding sites involved in DNA transcription, recombination and repair (38). We expect that new molecular insights into the interactions between IDRs and DNA sequences within the genomes of higher organisms may be obtainable using spectroscopic approaches that build on those of the present study.

## Protein Accession Code

The gp32 protein used in this work was prepared and purified as described in (31). The UniProt accession ID for this protein is: SSB_BPT4: P03965.

## Acknowledgements

The authors thank their laboratory colleagues for many helpful discussions. A.H.M. acknowledges Prof. Marina Guenza for helpful discussions regarding the gp32 protein. The work was supported by NIH NIGMS Grant GM-15792 to A.H.M. and P.H.v.H. P.H.v.H. is an American Cancer Society Research Professor of Chemistry.

## Supporting Information

### Sample preparation procedures

Fluorophore-labeled oligonucleotides were purchased from Integrated DNA Technologies (IDT, Coralville, IA, USA) and prepared in a standard aqueous buffer containing 100 mM NaCl, 6 mM MgCl_2_ (‘physiologically equivalent’ salt concentrations as discussed in the main text) and 10 mM Tris (pH 8), as described previously (1-3). Oxygen scavenging and triplet quenching reagents were used to extend the fluorescence lifetime and reduce the effects of triplet state blinking (4, 5). Samples were introduced into custom-built microfluidic sample chambers, which were constructed using fused silica quartz microscope slides. The inner surfaces of the slides were coated with a layer of biotinylated poly(ethylene glycol) and neutravidin. Individual DNA constructs were attached to the slide surfaces using the neutravidin linker, which binds tightly to the biotin attached to both the ss-dsDNA construct (see Table 1 of the main text) and to the poly(ethylene glycol) layer that coats the slide surface. The gp32 protein was prepared and purified as described in (3). Protein (gp32, UniProt SSB_BPT4: P03965) solutions at various concentrations were prepared in the standard aqueous buffer and introduced into the sample chamber for 2 – 3 minutes before data acquisition. Additional details of our sample preparation procedures are provided in (1-3).

### Microsecond-resolved single-molecule (sm) FRET experiments

The smFRET instrumentation and procedures used in these studies are identical to those described previously (1). Briefly, samples were illuminated using a 532 nm source laser in a total internal reflection fluorescence (TIRF) microscope. The laser beam was focused to a 50 *μ*m diameter spot and the incident power was adjusted to 10 mW. The Cy3 donor (*D*) and Cy5 acceptor (*A*) fluorescence from a single ss-dsDNA construct were collected using an oil-immersion objective (PlanApo, 100×, 1.4 NA, Nikon). Individual *D* and *A* photons were detected separately using avalanche photodiodes (APDs, SPCM-AQR-16, Perkin-Elmer) with 0.1 *μ*s resolution. The time-dependent fluorescence intensities were determined according to *I*_*D*(*A*)_(*t*) = *N*_*D*(*A*)_(*t*)/*T*_*w*_, where *N*_*D*(*A*)_ is the number of *D*(*A*) photons detected during an adjustable integration window, *T*_*w*_, centered at time *t*. The mean total fluorescence intensity, 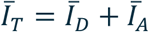, was typically ∼10,000 – 15,000 cps and the signal ‘background’ rate was ∼2,000 cps. Each single-molecule data set was recorded for a total duration time, *T* = ∼30 s. The ‘FRET efficiency’ was calculated following data acquisition as *E*^*FRET*^(*t*) = *I*_*A*_(*t*)/*I*_*T*_(*t*).

### Determination of probability distribution functions (PDFs) and two-point and three-point time correlation functions (TCFs)

The time-dependent FRET signal is related to the end-to-end *D*-*A* fluorophore distance, *R*_*DA*_, of the oligo(dT)_*n*_ template according to *E*^*FRET*^(*t*) = {1 + [*R*_*DA*_(*t*)/*R*_0_]^6^}^−1^, where *R*_0_ (≅ 56Å) is the Förster distance (6-8). From the smFRET trajectories, *E*^*FRET*^(*t*), we calculated the probability distribution functions (PDFs), *P*(*E*^*FRET*^), and the two-point and three-point time-correlation functions (TCFs), 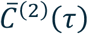 and 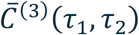, respectively (1, 2, 9-11). For a given trajectory of duration *T*, we divided the time series into *N* discrete intervals, *T*/*N* = Δ*t* = *T*_*w*_, such that the *i*th time interval was assigned to the integer index, *i* = *t*/Δ*t*. The time delay, *τ* = *n*Δ*t*, was thus assigned to the index *n*. We constructed the ensemble-averaged functions from ∼100 individual smFRET trajectories. The resulting functions from each trajectory were in turn averaged together.

We constructed the normalized PDF by summing over *N* discrete, time-averaged observations of microscopic configurations (termed ‘microstates’) according to

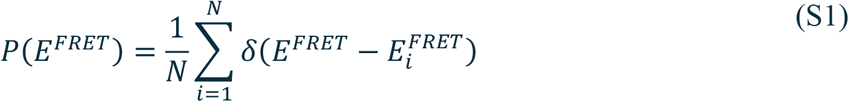

In Eq. (S1), *δ*(*x*) is the Dirac delta function and 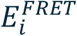 is the signal value determined for the *i*th time interval. The PDF contains information about the distribution of microstates at equilibrium averaged over the time scale of the adjustable integration window, *T*_*w*_. We calculated the normalized two-point TCF

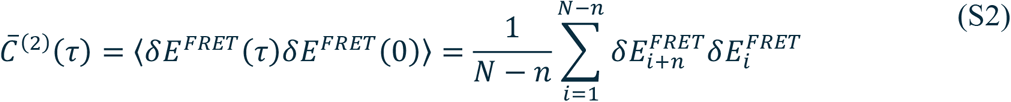

In Eq. (S2), 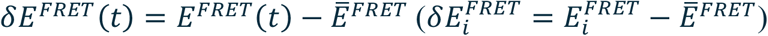 is the signal fluctuation about the mean value, 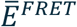. The two-point TCF contains information about transition pathways involving two successive observations that are separated by the interval, *τ*. We calculated the normalized three-point TCF according to

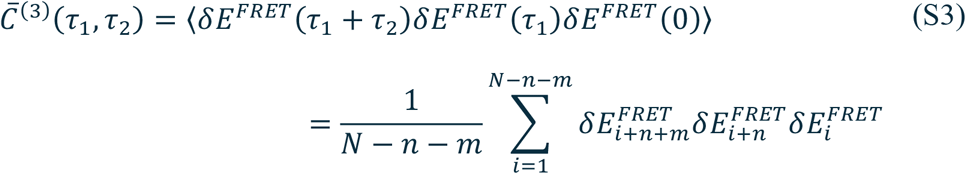

where the time intervals *τ*_1_ = *n*Δ*t* and *τ*_2_ = *m*Δ*t*. The three-point TCF contains information about transition pathways involving three successive observations separated by the intervals *τ*_1_ and *τ*_2_, and is thus sensitive to the presence of pathway intermediates that facilitate or hinder transitions between end-states (11).

### Determination of free energy surface (FES) parameters from kinetic network model analyses

As in previous work, we simulated the above functions using a kinetic network model (1, 2, 9, 11), which assumes that the Cy3/Cy5-oligo(dT)_*n*_-dsDNA / gp32 system interconverts between a finite number of Boltzmann-weighted microstates at equilibrium, and that the forward and backward rate constants are controlled by Arrhenius activation barriers. We note that individual microstates interconvert on time scales much faster than the microsecond resolution of our experiments. Thus, each ‘observation’ represents an average over the possible microstates that occur at a particular time, *t*, during the adjustable integration window, *T*_*w*_. It is useful to define a ‘macrostate’ as a collection of pseudo-degenerate microstates that occupy a common, locally stable region of the free energy surface (FES). We thus regard our microsecond-resolved single-molecule observations as direct observations of the dynamically interconverting macrostates of the system at equilibrium.

As we discuss in the following sections, for all our experiments the two-point TCFs can be described as a sum of four exponential decays with well-separated time constants. This implies that the system is composed of five dynamically interconverting macrostates at equilibrium (9). We model the time-dependent population of the *i*th macrostate, *p*_*i*_(*t*), using the master equation (12)

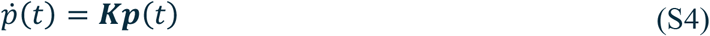

In Eq. (S4), ***p***(*t*) = [*p*_1_(*t*), *p*_2_(*t*), ⋯, *p*_5_(*t*)] is the time-dependent population vector and 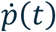 is its time derivative. The rate matrix ***K*** contains the forward (backward) rate constants *k*_*ij*(*ji*)_ between macrostates *i* and *j* [*i, j* ∈ {1,2,…,5}] according to

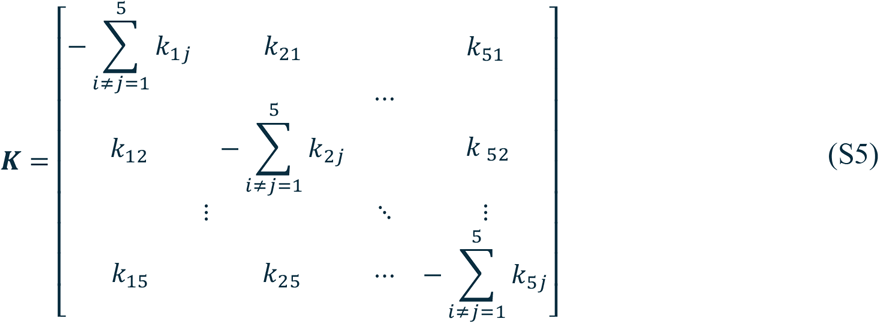

Given a set of input rate constants subject to completeness and detailed balance conditions (9), a solution to Eq. (S4) provides the probabilities of observing the *i*th macrostate at equilibrium, 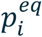, and the time-dependent conditional probabilities, *p*_*ij*_(*τ*), that the system undergoes a transition from macrostate *i* to macrostate *j* during the interval *τ*.

In our model the PDF is described as a sum of five macrostate contributions

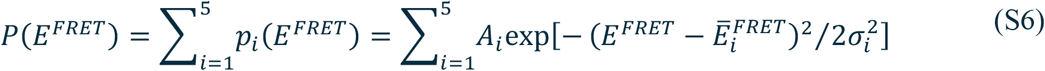

where we have assumed that the probability of the *i*th macrostate, *p*_*i*_(*E*^*FRET*^), has a Gaussian form with mean signal, 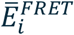, standard deviation *σ*_*i*_, and amplitude *A*_*i*_. The probability that the *i*th macrostate is present at equilibrium is the integrated area 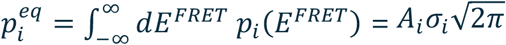.

The two-point TCF can be described according to

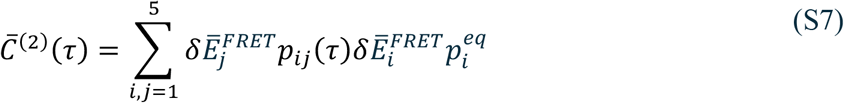

where 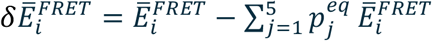 is the signal fluctuation of the *i*th macrostate. According to Eq. (S6), 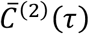 is a sum of 5 × 5 = 25 ‘kinetic pathways’ in which two consecutive fluctuations of the signal occur, first with 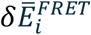 at *τ* = 0 and second with 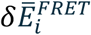 at time *τ*. Since the signs of the fluctuations 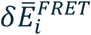 can be positive or negative, the various two-point products, 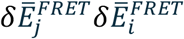 contribute both positive and negative amplitudes to the overall sum. The statistical weights of the various pathways depend on the products of probabilities, 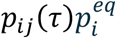, which depend on *τ*. Thus, the two-point TCF is the statistically-weighted product of two consecutive fluctuations, which occur within the interval *τ* as the system undergoes spontaneous transitions between the five macrostates.

We simulated the three-point TCF according to

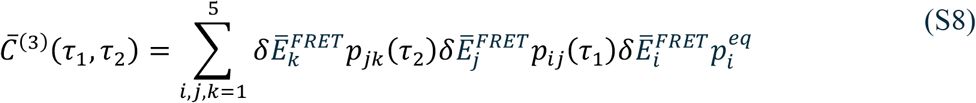

Equation (S8) shows that the function 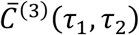 is a sum of (5)^3^ = 125 ‘transition pathways’ in which three consecutive fluctuations of the visibility occur – initially with 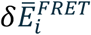, and then with 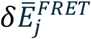 at time *τ*_1_ and 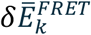 at time *τ*_2_. The various three-point products, 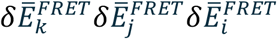 contribute both positive and negative amplitudes to the overall sum, with relative weights given by the product of probabilities, 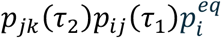. We thus view the three-point TCF as the statistically-weighted product of three consecutive fluctuations separated by the intervals *τ*_1_ and *τ*_2_. The three-point TCF is sensitive to the roles of transient intermediates in the conformational transition pathways that connects the five macrostates at equilibrium.

As an initial step towards simulating the statistical functions described above, it was necessary to determine the minimum number of macrostates, *M*, that can represent the data. The general solution to Eq. (S4) requires that the two-point TCF is the sum of *M* − 1 decaying exponentials (9, 12). For all the samples that we studied the two-point TCFs are well described as the sum of five exponential decays with time constants separated by at least one order of magnitude (see Fig. S1 and Table S1). We determined that the decay component with the slowest time constant (∼ 1 s) is due to mechanical laboratory noise, as shown in previous work (1, 10). We therefore assume a kinetic network model with *M* = 5.

We performed a multiparameter variational calculation to find ‘optimized’ solutions of Eq. (4) that best describe our smFRET data. For each of the 3’,5’-Cy3/Cy5-oligo(dT)_14,15_-dsDNA constructs discussed in the main text, we calculated the statistical functions, *P*(*E*^*FRET*^), 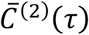 and 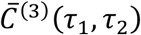, using Eqs. (S6) – (S8), respectively, for a given set of input rate constants, *k*_*ij*_, mean FRET values, 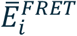, amplitudes, *A*_*i*_, and standard deviations, *σ*_*i*_. Our analysis involved determining the connectivity between macrostates as defined by the elements of the rate matrix ***K*** [Eq. (S5)]. Because the parameter space of possible five-state models is large, we developed and implemented an algorithm to generate and test all possible models in parallel, as described in recent work (11). We thus efficiently explored the parameter space and tested hundreds of different network configurations in parallel using a high-performance computer cluster to arrive at the final optimized solutions.

To constrain the number of possible network configurations, we imposed the following assumptions based on our experience from previous studies (1-3, 9). We assigned the *S*_1_ macrostate (with the highest FRET value, 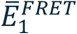) to conformations in which the 3’,5’-Cy3/Cy5-oligo(dT)_14,15_-dsDNA constructs are not bound to protein (see, *e*.*g*., Fig. 1*B* of the main text). Similarly, we assigned the *S*_5_ macrostate (with the lowest FRET value, 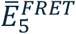) to conformations in which the oligo(dT)_14,15_ region of the DNA construct is maximally extended due to the presence of two cooperatively-bound gp32 proteins (*i*.*e*., a dimer-bound state, as shown in Fig. 1*D*). Previous studies have established that gp32 proteins bind initially as monomers to ssDNA templates in a noncooperative fashion. In a subsequent step the 3’,5’-Cy3/Cy5-oligo(dT)_14,15_-dsDNA-gp32 monomer complex can associate with a second gp32 protein to form a cooperatively-bound 3’,5’-Cy3/Cy5-oligo(dT)_14,15_-dsDNA-(gp32)_2_ dimer complex (2, 3). We thus assigned the three macrostates with intermediate FRET values, labeled *S*_2_ − *S*_4_, to distinct monomer-bound complexes.

We next consider the possible pathways in which a noncooperatively-bound monomer can function as a reactive intermediate connecting the end-states, *S*_1_ and *S*_5_. In one pathway, a monomer-bound complex can form by association close to the ss-dsDNA junction (at lattice sites 1 – 7 or 2 – 8, as depicted in Fig. 1*C*) without occluding the 7 contiguous lattice sites required for a second protein to form the dimer state, *S*_5_ (depicted in Fig. 1*D*). In a second pathway, a monomer-bound complex can form by association close to the nascent end of the oligo(dT)_14,15_ region of the DNA construct (at lattice sites 8 – 14 or 9 – 15), which would also provide a contiguous set of *s* = 7 vacant lattice sites to permit a second protein to form the dimer state, *S*_5_. In both cases, we refer to the monomer-bound states as ‘productive’ intermediates’ because they can lead to the formation of the dimer-DNA-bound complex. In a third pathway, the protein binds at a lattice position near the middle of the oligo(dT)_14,15_ region of the DNA construct, thereby occluding adjacent lattice sites and preventing a second gp32 protein from immediately binding and forming the dimer-DNA-bound complex. Under conditions in which this latter monomer-DNA-bound state is unstable, or has a short lifetime, it represents a kinetic ‘dead-end’ on the assembly pathway of the dimer-DNA-bound complex.

We quantified the agreement between calculated and experimentally-derived functions using a nonlinear least squares target function, χ^2^. We thus obtained ‘optimized’ solutions to the master equation by performing an iterative search of the parameter space to minimize the target function, following a procedure like that described in previous studies (1, 2, 9, 11). From these calculations, we determined optimized values of the forward and backward time constants, 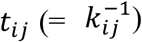, and the equilibrium populations, 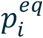. Using these parameters we determined the Boltzmann-weighted relative stabilities 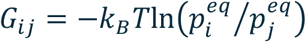, and the relative Arrhenius activation barriers, 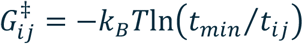, where *k*_*B*_ is Boltzmann’s constant and *t*_*min*_ is a reference time scale equal to that of the fastest process measured in the experiment (typically sub-millisecond).

**Figure S1.**
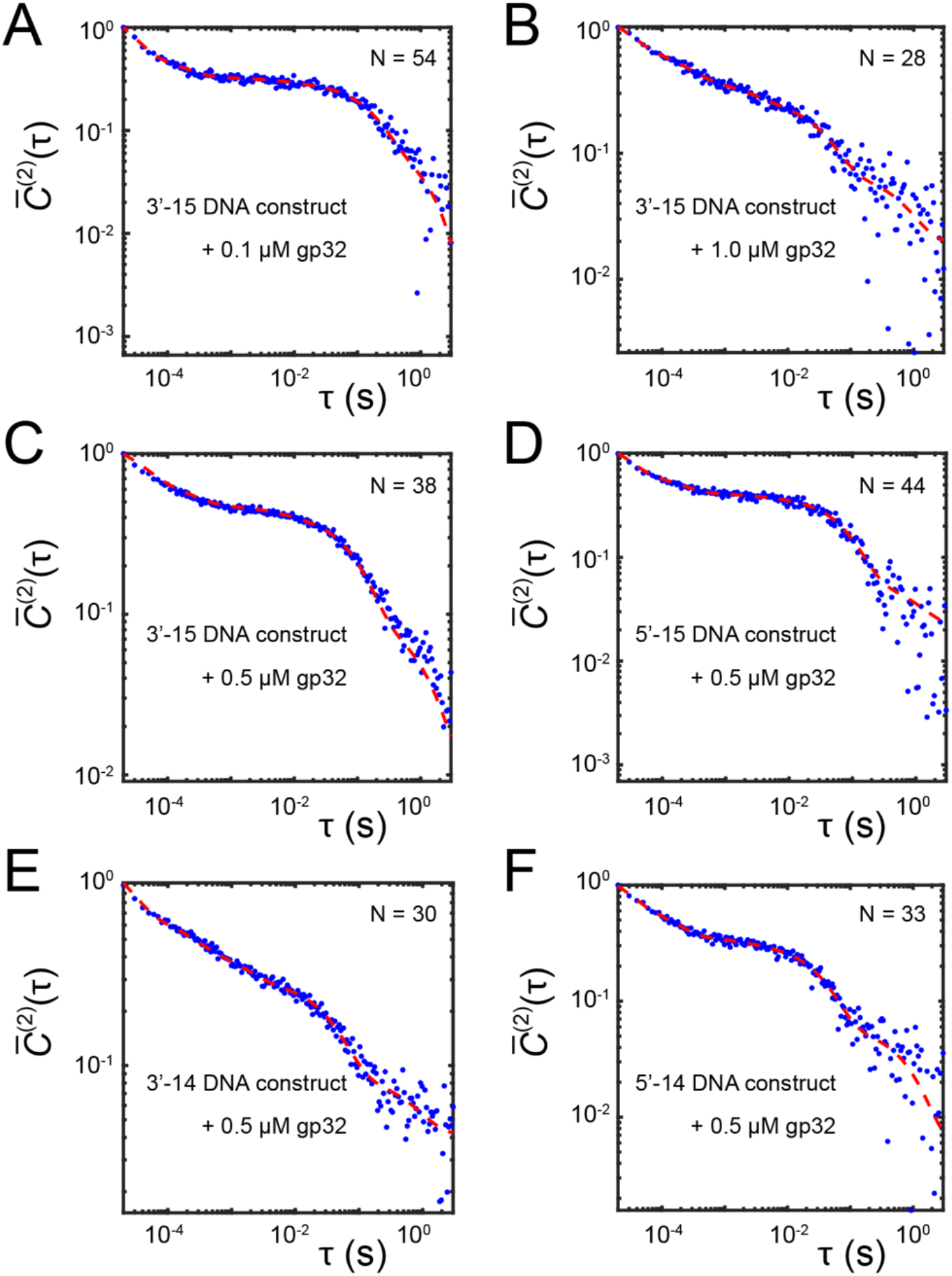
Two-point TCFs of the DNA constructs used in this work in the presence of gp32 protein. The base sequences of the DNA constructs are listed in Table 1 of the main text. Data are shown as blue points. Red curves are the model five-exponential functions: 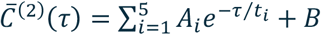, with parameters listed in Table S1. The number of data sets, N, that were used to construct each TCF are indicated. (***A***) 3’-15 DNA construct in 0.1 *μ*M gp32. (***B***) 3’-15 DNA construct in 1.0 *μ*M gp32. (***C***) 3’-15 DNA construct in 0.5 *μ*M gp32. (***D***) 5’-15 DNA construct in 0.5 *μ*M gp32. (***E***) 3’-14 DNA construct in 0.5 *μ*M gp32. (***F***) 5’-14 DNA construct in 0.5 *μ*M gp32. Experiments were performed at room temperature (23 °C) and physiological buffer salt conditions ([NaCl] = 100 mM, [MgCl_2_] = 6 mM and 10 mM Tris, pH 8.0).

**Table S1.** Optimized parameters for the model five-exponential functions 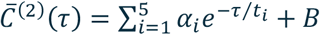. *R*^2^ is the goodness-of-fit parameter. Time constants are given in milliseconds. The slowest decay component, *t*_5_ (∼1 s), is due to mechanical laboratory noise. Fits correspond to the red curves shown in Fig. S1.

| Sample | $\alpha_1$ | $t_1$ | $\alpha_2$ | $t_2$ | $\alpha_3$ | $t_3$ | $\alpha_4$ | $t_4$ | $\alpha_5$ | $t_5$ | $B$ | $R^2$ |
| --- | --- | --- | --- | --- | --- | --- | --- | --- | --- | --- | --- | --- |
| 3'-15,<br>0.1 μM | 1.26 | 0.020 | 0.23 | 0.17 | 0.03 | 6.8 | 0.22 | 167 | 0.08 | 1340 | 0.02 | 0.992 |
| 3'-15,<br>1.0 μM | 0.78 | 0.024 | 0.32 | 0.22 | 0.12 | 2.5 | 0.20 | 38.2 | 0.05 | 710 | 0.02 | 0.988 |
| 3'-15,<br>0.5 μM | 0.56 | 0.034 | 0.22 | 0.22 | 0.06 | 5.2 | 0.33 | 98.5 | 0.08 | 1230 | 0.00 | 0.997 |
| 5'-15,<br>0.5 μM | 0.92 | 0.020 | 0.26 | 0.17 | 0.03 | 2.8 | 0.33 | 77.5 | 0.04 | 946 | 0.02 | 0.988 |
| 3'-14,<br>0.5 μM | 0.95 | 0.020 | 0.25 | 0.20 | 0.16 | 1.5 | 0.21 | 42.4 | 0.05 | 747 | 0.04 | 0.994 |
| 5'-14,<br>0.5 μM | 0.86 | 0.020 | 0.36 | 0.14 | 0.04 | 2.1 | 0.26 | 36.4 | 0.05 | 868 | 0.01 | 0.990 |

**Figure S2.**
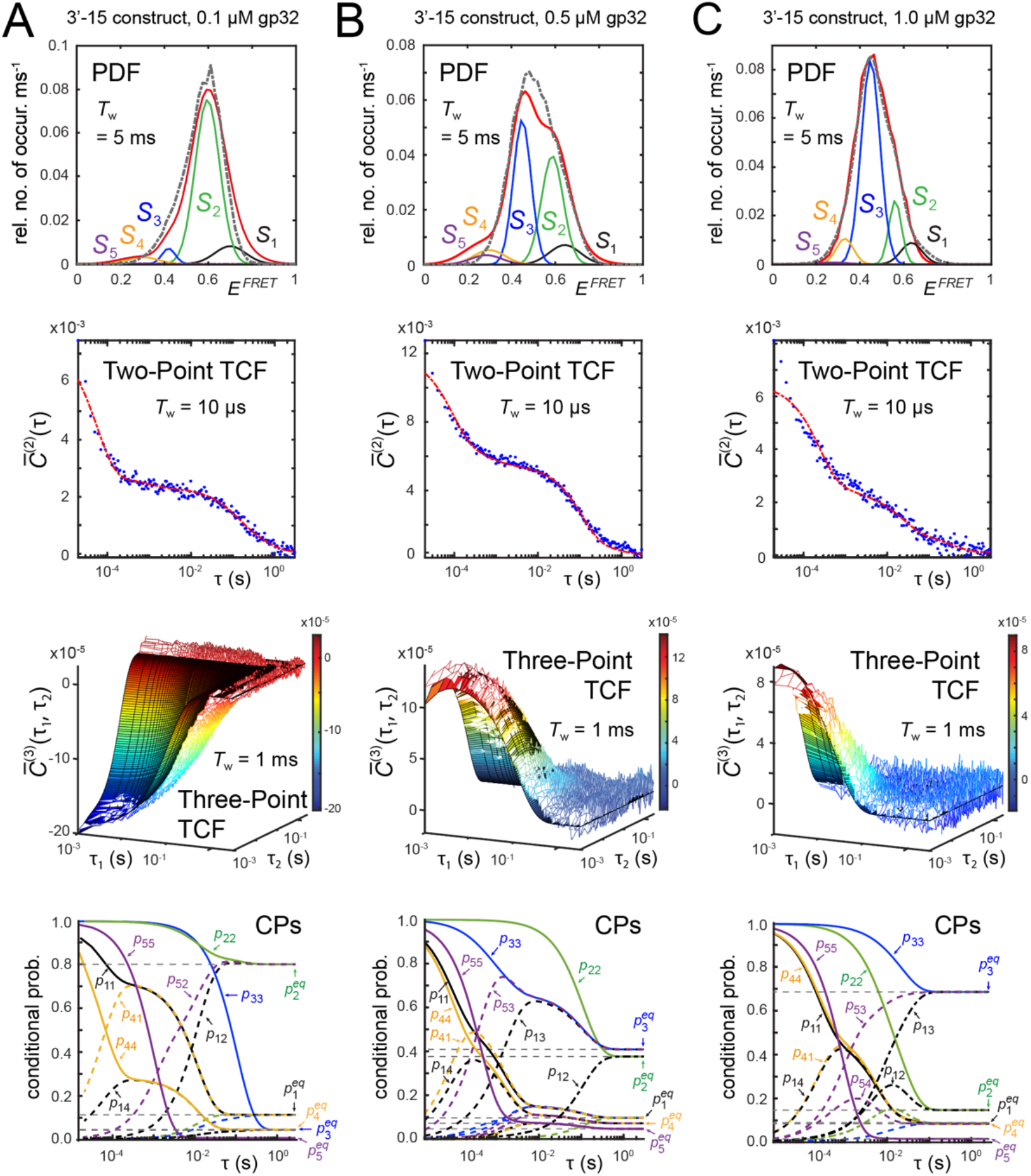
Results of the five-state kinetic network analysis applied to microsecond-resolved smFRET measurements of the 3’-Cy3/Cy5-oligo(dT)_15_-dsDNA construct in 10 mM Tris at pH 8.0, 100 mM NaCl, 6 mM MgCl_2_ and (***A***) 0.1 μM gp32; (***B***) 0.5 μM gp32; (***C***) 1.0 μM gp32. Optimized kinetic and equilibrium parameters are obtained from simultaneously fitting experimental PDFs (black dashed curves, first row), the two-point TCFs (blue dots, second row), the three-point TCFs (colored mesh, third row) and the CPs [*p*_*ij*_(*τ*) with *i, j* ∈ {1,2,3,4,5}, fourth row]. Integration periods are indicated in the insets. The model PDFs (red curves) are modeled as sums of five Gaussian macrostates (black, green, blue, orange, and purple curves, labeled *S*_1_ – *S*_5_, respectively) with areas equal to the equilibrium probabilities, 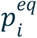. The model two-point TCFs are shown as red dashed curves, and the model three-point TCFs are shown as solid surfaces. The CPs are shown as solid curves for *i* = *j* (called ‘survival probabilities’), and as dashed curves for *i* ≠ *j* (‘transition probabilities’). The CPs are color coded as the state labels, according to the value of *i* = 1: black, 2: green, 3: blue, 4: orange and 5: purple. CPs are shown that contribute to the kinetic network scheme.

**Table S2.** Optimized parameters for the Gaussian components of the microsecond-resolved smFRET probability distribution functions (PDFs).

| Sample | $\bar{E}_1^{FRET}$ | $\bar{E}_2^{FRET}$ | $\bar{E}_3^{FRET}$ | $\bar{E}_4^{FRET}$ | $\bar{E}_5^{FRET}$ | $\sigma_1$ | $\sigma_2$ | $\sigma_3$ | $\sigma_4$ | $\sigma_5$ | $A_1$ | $A_2$ | $A_3$ | $A_4$ | $A_5$ |
| --- | --- | --- | --- | --- | --- | --- | --- | --- | --- | --- | --- | --- | --- | --- | --- |
| 3'-15,<br>0.1 μM | 0.70 | 0.59 | 0.42 | 0.30 | 0.29 | 0.10 | 0.08 | 0.04 | 0.10 | 0.10 | 0.44 | 3.98 | 0.40 | 0.16 | 0.04 |
| 3'-15,<br>1.0 μM | 0.63 | 0.56 | 0.45 | 0.33 | 0.27 | 0.07 | 0.04 | 0.06 | 0.06 | 0.10 | 0.46 | 1.39 | 4.58 | 0.53 | 0.04 |
| 3'-15,<br>0.5 μM | 0.64 | 0.59 | 0.45 | 0.30 | 0.29 | 0.10 | 0.07 | 0.06 | 0.10 | 0.10 | 0.40 | 2.11 | 2.72 | 0.28 | 0.24 |
| 5'-15,<br>0.5 μM | 0.70 | 0.60 | 0.48 | 0.30 | 0.20 | 0.08 | 0.05 | 0.08 | 0.10 | 0.10 | 1.99 | 1.04 | 1.99 | 0.24 | 0.04 |
| 3'-14,<br>0.5 μM | 0.66 | 0.58 | 0.47 | 0.30 | 0.23 | 0.05 | 0.05 | 0.07 | 0.09 | 0.10 | 1.12 | 1.51 | 3.13 | 0.26 | 0.24 |
| 5'-14,<br>0.5 μM | 0.70 | 0.60 | 0.48 | 0.30 | 0.20 | 0.06 | 0.06 | 0.07 | 0.10 | 0.10 | 1.99 | 2.19 | 1.71 | 0.24 | 0.04 |

Parameters defined in Eq. (5). Error bars for the mean FRET efficiencies, $\bar{E}_i^{FRET}$ , are <1% of reported values. PDFs were constructed using the integration period $T_w = 5$ ms. All samples prepared in 10 mM Tris at pH 8.0 and 100 mM NaCl, 6 mM MgCl<sub>2</sub>.

**Table S3.**
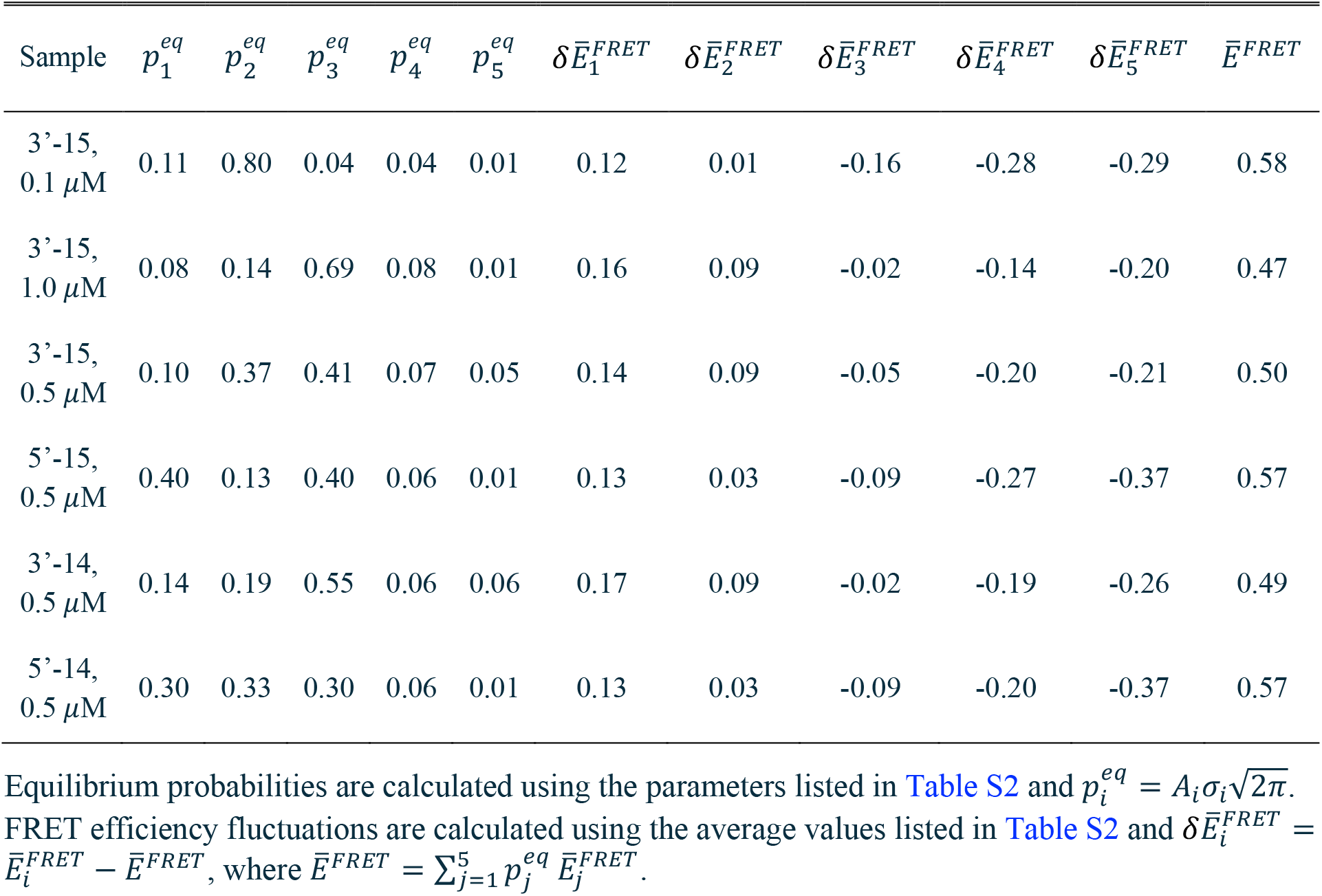
Optimized values of the equilibrium probabilities and signal fluctuations.

| Sample | $p_1^{eq}$ | $p_2^{eq}$ | $p_3^{eq}$ | $p_4^{eq}$ | $p_5^{eq}$ | $\delta\bar{E}_1^{FRET}$ | $\delta\bar{E}_2^{FRET}$ | $\delta\bar{E}_3^{FRET}$ | $\delta\bar{E}_4^{FRET}$ | $\delta\bar{E}_5^{FRET}$ | $\bar{E}^{FRET}$ |
| --- | --- | --- | --- | --- | --- | --- | --- | --- | --- | --- | --- |
| 3'-15,<br>0.1 μM | 0.11 | 0.80 | 0.04 | 0.04 | 0.01 | 0.12 | 0.01 | -0.16 | -0.28 | -0.29 | 0.58 |
| 3'-15,<br>1.0 μM | 0.08 | 0.14 | 0.69 | 0.08 | 0.01 | 0.16 | 0.09 | -0.02 | -0.14 | -0.20 | 0.47 |
| 3'-15,<br>0.5 μM | 0.10 | 0.37 | 0.41 | 0.07 | 0.05 | 0.14 | 0.09 | -0.05 | -0.20 | -0.21 | 0.50 |
| 5'-15,<br>0.5 μM | 0.40 | 0.13 | 0.40 | 0.06 | 0.01 | 0.13 | 0.03 | -0.09 | -0.27 | -0.37 | 0.57 |
| 3'-14,<br>0.5 μM | 0.14 | 0.19 | 0.55 | 0.06 | 0.06 | 0.17 | 0.09 | -0.02 | -0.19 | -0.26 | 0.49 |
| 5'-14,<br>0.5 μM | 0.30 | 0.33 | 0.30 | 0.06 | 0.01 | 0.13 | 0.03 | -0.09 | -0.20 | -0.37 | 0.57 |

**Table S4.** Optimized forward and backward inverse rate constants corresponding to the elementary reactive steps for the samples studied.

| Sample | $k_{12}^{-1}$ | $k_{21}^{-1}$ | $k_{13}^{-1}$ | $k_{31}^{-1}$ | $k_{14}^{-1}$ | $k_{41}^{-1}$ | $k_{25}^{-1}$ | $k_{52}^{-1}$ | $k_{45}^{-1}$ | $k_{54}^{-1}$ | $k_{35}^{-1}$ | $k_{53}^{-1}$ |
| --- | --- | --- | --- | --- | --- | --- | --- | --- | --- | --- | --- | --- |
| 3'-15, 0.1 $\mu\text{M}$ | 10.9 | 93.0 | 300 | 112 | 0.20 | 0.10 | 258 | 2.49 | 57.9 | 1.82 | - | - |
| 3'-15, 1.0 $\mu\text{M}$ | 8.40 | 14.6 | 9.65 | 81.9 | 0.51 | 0.10 | - | - | 16.4 | 1.97 | 123 | 1.80 |
| 3'-15 0.5 $\mu\text{M}$ | 44.7 | 174 | 1.27 | 5.50 | 0.21 | 0.10 | - | - | 30.8 | 11.5 | 4.09 | 0.48 |
| 5'-15, 0.5 $\mu\text{M}$ | 120 | 39.0 | 132 | 300 | 0.69 | 0.10 | - | - | 13.7 | 1.10 | 232 | 51.6 |
| 3'-14, 0.5 $\mu\text{M}$ | 32.4 | 44.7 | 7.25 | 45.5 | 0.18 | 0.10 | - | - | 20.8 | 6.16 | 3.83 | 0.45 |
| 5'-14, 0.5 $\mu\text{M}$ | 15.4 | 17.3 | 48.1 | 89.2 | 0.69 | 0.10 | - | - | 34.4 | 1.54 | 88.4 | 10.4 |

Inverse rate constants defined in Eq. (4), listed in units of milliseconds. Error bars are < 1% of reported values (see Statistical Uncertainty Analysis below).

**Table S5.** Relative free energy minima.

| Sample | $G_1$ | $G_2$ | $G_3$ | $G_4$ | $G_5$ |
| --- | --- | --- | --- | --- | --- |
| 3'-15, 0.1 $\mu\text{M}$ | 2.22 | 0 | 2.36 | 3.18 | 5.53 |
| 3'-15, 1.0 $\mu\text{M}$ | 2.25 | 1.16 | 0 | 2.07 | 4.75 |
| 3'-15 0.5 $\mu\text{M}$ | 2.00 | 0.29 | 0 | 2.30 | 2.73 |
| 5'-15, 0.5 $\mu\text{M}$ | 0.05 | 0.68 | 0 | 2.17 | 4.45 |
| 3'-14, 0.5 $\mu\text{M}$ | 1.03 | 0.57 | 0 | 2.42 | 2.50 |
| 5'-14, 0.5 $\mu\text{M}$ | 0.21 | 0 | 0.33 | 2.26 | 4.43 |

**Table S6.**
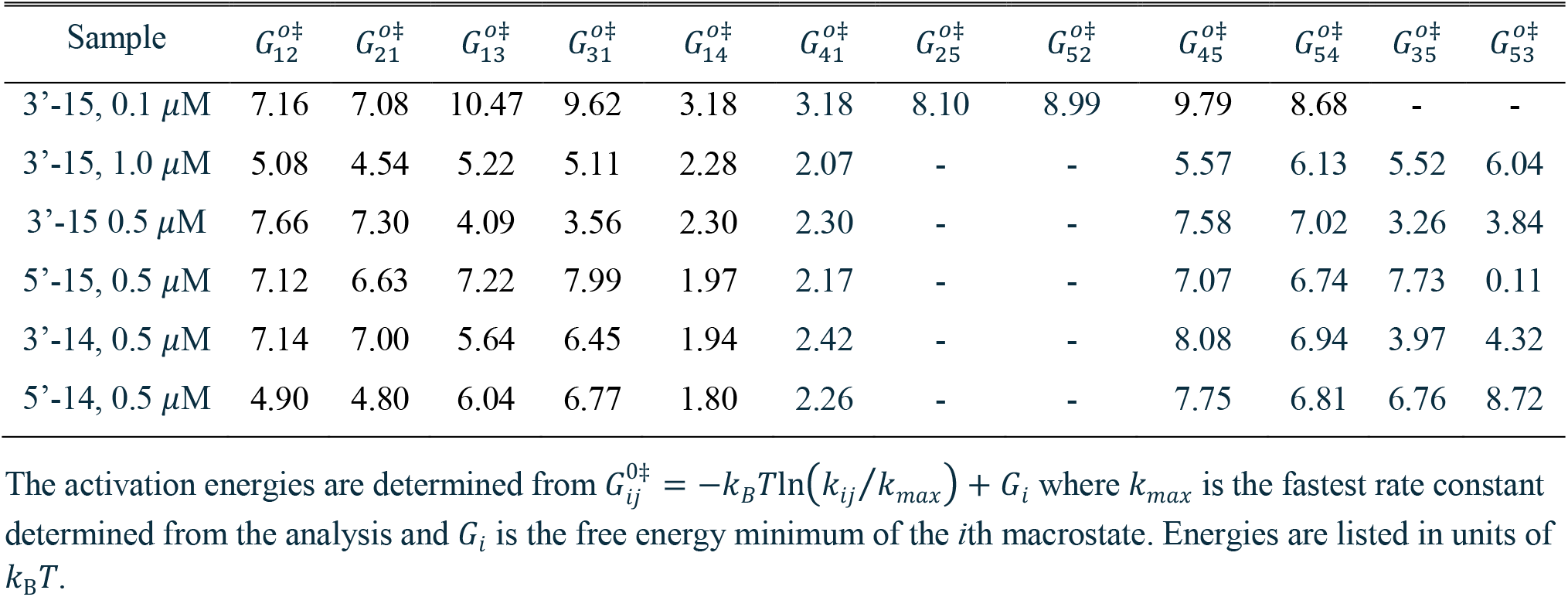
Free energies of activation corresponding to the elementary reactive steps.

**Figure S3.**
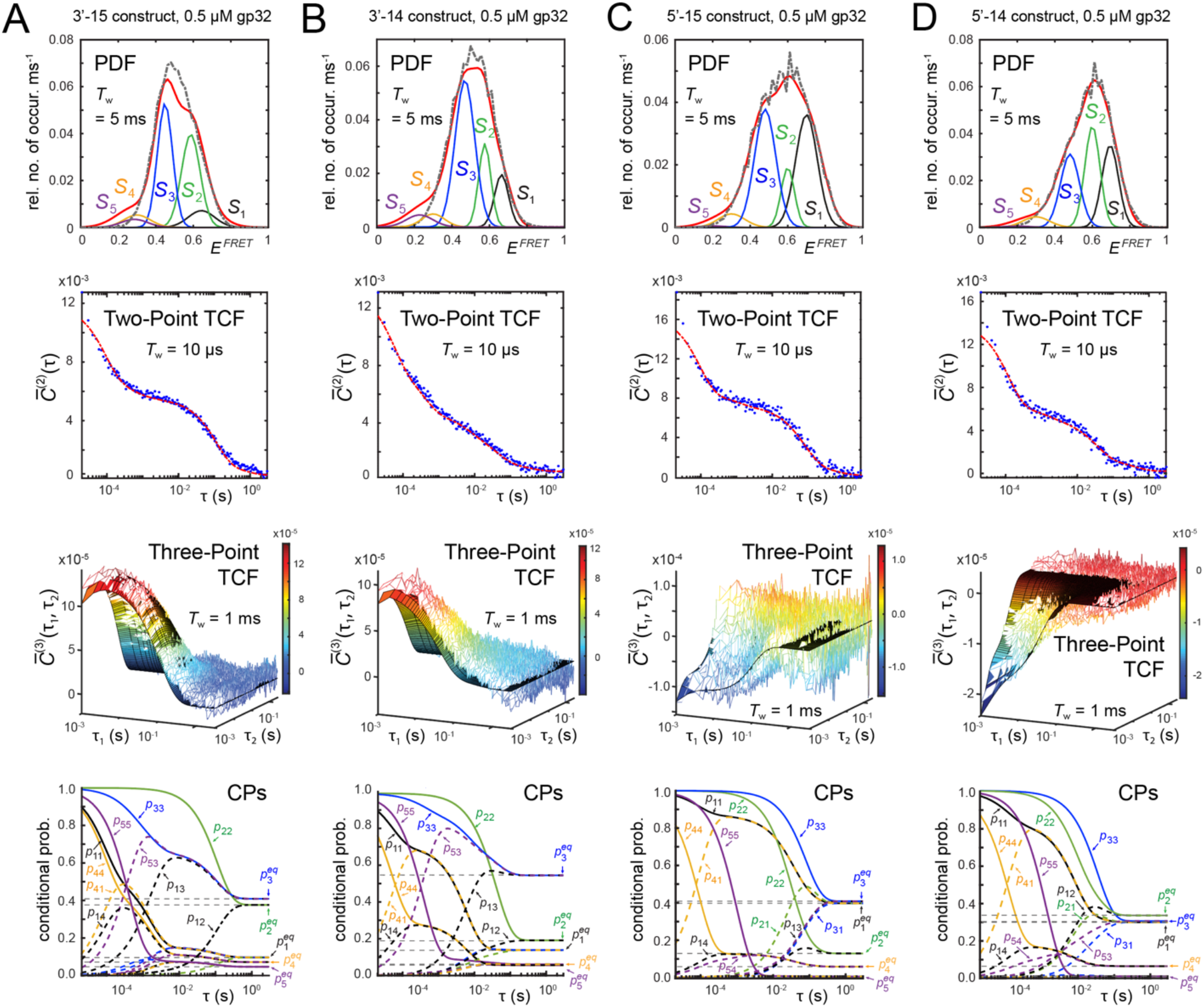
Results of five-state kinetic network model analysis applied to microsecond-resolved smFRET measurements of the (***A***) 3’-15; (***B***) 3’-14; (***C***) 5’-15 and (***D***) 5’-14 DNA constructs (see Table 1 of the main text) in 10 mM Tris at pH 8.0, 100 mM NaCl and 6 mM MgCl_2_. Formatting and color schemes are the same as described in Fig. S2.

**Table S7.** Decomposition of the largest pathway terms for the two-point TCFs [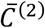 function, Eq. (7) of main text] at *τ* = 10 μs.

| 3'-15 DNA construct + 0.1 $\mu\text{M}$ gp32 | | | | | | | | | |
| --- | --- | --- | --- | --- | --- | --- | --- | --- | --- |
| Positive terms |  |  |  |  | Negative terms |  |  |  |  |
| $i$ | $j$ | $\delta\bar{E}_i\delta\bar{E}_j$ | $w_{ij}$ | $w_{ij}\delta\bar{E}_i\delta\bar{E}_j$ | $i$ | $j$ | $\delta\bar{E}_i\delta\bar{E}_j$ | $w_{ij}$ | $w_{ij}\delta\bar{E}_i\delta\bar{E}_j$ |
| 4 | 4 | $8.22 \times 10^{-2}$ | $3.32 \times 10^{-2}$ | $2.73 \times 10^{-3}$ | 4 | 1 | $-3.25 \times 10^{-2}$ | $9.11 \times 10^{-3}$ | $-2.96 \times 10^{-4}$ |
| 1 | 1 | $1.28 \times 10^{-2}$ | $1.01 \times 10^{-1}$ | $1.29 \times 10^{-3}$ | 1 | 4 | $-3.25 \times 10^{-2}$ | $9.08 \times 10^{-3}$ | $-2.95 \times 10^{-4}$ |
| 3 | 3 | $2.81 \times 10^{-2}$ | $4.10 \times 10^{-2}$ | $1.16 \times 10^{-3}$ | 2 | 5 | $-2.79 \times 10^{-3}$ | $6.16 \times 10^{-5}$ | $-1.72 \times 10^{-7}$ |
| 5 | 5 | $8.57 \times 10^{-2}$ | $3.96 \times 10^{-3}$ | $3.39 \times 10^{-4}$ | 5 | 1 | $-3.31 \times 10^{-2}$ | $5.03 \times 10^{-6}$ | $-1.67 \times 10^{-7}$ |
| 2 | 2 | $9.11 \times 10^{-5}$ | $8.02 \times 10^{-1}$ | $7.31 \times 10^{-5}$ | 3 | 1 | $-1.90 \times 10^{-2}$ | $7.05 \times 10^{-6}$ | $-1.34 \times 10^{-7}$ |
| 5 | 4 | $8.39 \times 10^{-2}$ | $3.89 \times 10^{-5}$ | $3.27 \times 10^{-6}$ | 1 | 3 | $-1.90 \times 10^{-2}$ | $7.03 \times 10^{-6}$ | $-1.33 \times 10^{-7}$ |
| 4 | 5 | $8.39 \times 10^{-2}$ | $1.28 \times 10^{-5}$ | $1.08 \times 10^{-6}$ | 5 | 2 | $-2.79 \times 10^{-3}$ | $3.22 \times 10^{-5}$ | $-8.99 \times 10^{-8}$ |
| 1 | 2 | $1.08 \times 10^{-3}$ | $1.93 \times 10^{-4}$ | $2.09 \times 10^{-7}$ | 1 | 5 | $-3.31 \times 10^{-2}$ | $1.66 \times 10^{-6}$ | $-5.49 \times 10^{-8}$ |
| 2 | 1 | $1.08 \times 10^{-3}$ | $1.65 \times 10^{-4}$ | $1.78 \times 10^{-7}$ | 4 | 2 | $-2.74 \times 10^{-3}$ | $8.89 \times 10^{-6}$ | $-2.43 \times 10^{-8}$ |
| 3 | 4 | $4.81 \times 10^{-2}$ | $3.22 \times 10^{-7}$ | $1.55 \times 10^{-8}$ | 2 | 4 | $-2.74 \times 10^{-3}$ | $7.83 \times 10^{-6}$ | $-2.14 \times 10^{-8}$ |

| 3'-15 DNA construct + 1.0 $\mu\text{M}$ gp32 | | | | | | | | | |
| --- | --- | --- | --- | --- | --- | --- | --- | --- | --- |
| Positive terms |  |  |  |  | Negative terms |  |  |  |  |
| $i$ | $j$ | $\delta\bar{E}_i\delta\bar{E}_j$ | $w_{ij}$ | $w_{ij}\delta\bar{E}_i\delta\bar{E}_j$ | $i$ | $j$ | $\delta\bar{E}_i\delta\bar{E}_j$ | $w_{ij}$ | $w_{ij}\delta\bar{E}_i\delta\bar{E}_j$ |
| 1 | 1 | $2.77 \times 10^{-2}$ | $7.85 \times 10^{-2}$ | $2.17 \times 10^{-3}$ | 4 | 1 | $-2.30 \times 10^{-2}$ | $3.07 \times 10^{-3}$ | $-7.10 \times 10^{-5}$ |
| 4 | 4 | $1.91 \times 10^{-2}$ | $7.64 \times 10^{-2}$ | $1.46 \times 10^{-3}$ | 1 | 4 | $-2.30 \times 10^{-2}$ | $3.07 \times 10^{-3}$ | $-7.00 \times 10^{-5}$ |
| 2 | 2 | $8.46 \times 10^{-3}$ | $1.42 \times 10^{-1}$ | $1.20 \times 10^{-3}$ | 1 | 3 | $-3.36 \times 10^{-3}$ | $1.66 \times 10^{-4}$ | $-5.60 \times 10^{-7}$ |
| 5 | 5 | $3.76 \times 10^{-2}$ | $9.60 \times 10^{-3}$ | $3.61 \times 10^{-3}$ | 3 | 1 | $-3.36 \times 10^{-3}$ | $1.64 \times 10^{-4}$ | $-5.50 \times 10^{-7}$ |
| 3 | 3 | $4.07 \times 10^{-4}$ | $6.86 \times 10^{-1}$ | $2.79 \times 10^{-4}$ | 5 | 1 | $-3.23 \times 10^{-2}$ | $1.95 \times 10^{-6}$ | $-6.30 \times 10^{-8}$ |
| 2 | 1 | $1.53 \times 10^{-2}$ | $1.91 \times 10^{-4}$ | $2.92 \times 10^{-6}$ | 1 | 5 | $-3.23 \times 10^{-2}$ | $1.90 \times 10^{-6}$ | $-6.10 \times 10^{-8}$ |
| 1 | 2 | $1.53 \times 10^{-2}$ | $1.91 \times 10^{-4}$ | $2.92 \times 10^{-6}$ | 4 | 2 | $-1.27 \times 10^{-2}$ | $3.71 \times 10^{-6}$ | $-4.70 \times 10^{-8}$ |
| 5 | 4 | $2.68 \times 10^{-2}$ | $9.65 \times 10^{-5}$ | $2.58 \times 10^{-6}$ | 2 | 4 | $-1.27 \times 10^{-2}$ | $3.70 \times 10^{-6}$ | $-4.70 \times 10^{-8}$ |
| 4 | 5 | $2.68 \times 10^{-2}$ | $9.42 \times 10^{-5}$ | $2.52 \times 10^{-6}$ | 2 | 3 | $-1.86 \times 10^{-3}$ | $1.99 \times 10^{-7}$ | $-3.70 \times 10^{-10}$ |
| 3 | 5 | $3.92 \times 10^{-3}$ | $1.10 \times 10^{-4}$ | $4.30 \times 10^{-7}$ | 3 | 2 | $-1.86 \times 10^{-3}$ | $1.96 \times 10^{-7}$ | $-3.60 \times 10^{-10}$ |

| 3'-15 DNA construct + 0.5 μM gp32 | | | | | | | | | |
| --- | --- | --- | --- | --- | --- | --- | --- | --- | --- |
| Positive terms |  |  |  |  | Negative terms |  |  |  |  |
| $i$ | $j$ | $\delta \bar{E}_i \delta \bar{E}_j$ | $w_{ij}$ | $w_{ij} \delta \bar{E}_i \delta \bar{E}_j$ | $i$ | $j$ | $\delta \bar{E}_i \delta \bar{E}_j$ | $w_{ij}$ | $w_{ij} \delta \bar{E}_i \delta \bar{E}_j$ |
| 2 | 2 | $7.26 \times 10^{-3}$ | $3.76 \times 10^{-1}$ | $2.73 \times 10^{-3}$ | 4 | 1 | $-2.86 \times 10^{-2}$ | $8.15 \times 10^{-3}$ | $-2.33 \times 10^{-4}$ |
| 4 | 4 | $4.06 \times 10^{-2}$ | $6.35 \times 10^{-2}$ | $2.57 \times 10^{-3}$ | 1 | 4 | $-2.86 \times 10^{-2}$ | $8.12 \times 10^{-3}$ | $-2.32 \times 10^{-4}$ |
| 5 | 5 | $4.64 \times 10^{-2}$ | $4.46 \times 10^{-2}$ | $2.07 \times 10^{-3}$ | 1 | 3 | $-7.40 \times 10^{-3}$ | $1.44 \times 10^{-3}$ | $-1.06 \times 10^{-5}$ |
| 1 | 1 | $2.02 \times 10^{-2}$ | $8.70 \times 10^{-2}$ | $1.75 \times 10^{-3}$ | 3 | 1 | $-7.41 \times 10^{-3}$ | $1.40 \times 10^{-3}$ | $-1.04 \times 10^{-5}$ |
| 3 | 3 | $2.72 \times 10^{-3}$ | $4.06 \times 10^{-1}$ | $1.10 \times 10^{-3}$ | 5 | 1 | $-3.06 \times 10^{-2}$ | $8.13 \times 10^{-6}$ | $-2.49 \times 10^{-7}$ |
| 3 | 5 | $1.12 \times 10^{-2}$ | $1.95 \times 10^{-3}$ | $2.19 \times 10^{-5}$ | 1 | 5 | $-3.06 \times 10^{-2}$ | $6.23 \times 10^{-6}$ | $-1.91 \times 10^{-7}$ |
| 5 | 3 | $1.12 \times 10^{-2}$ | $1.91 \times 10^{-3}$ | $2.15 \times 10^{-5}$ | 4 | 2 | $-1.72 \times 10^{-2}$ | $1.90 \times 10^{-6}$ | $-3.26 \times 10^{-8}$ |
| 5 | 4 | $4.34 \times 10^{-2}$ | $7.50 \times 10^{-5}$ | $3.25 \times 10^{-6}$ | 2 | 4 | $-1.72 \times 10^{-2}$ | $1.89 \times 10^{-6}$ | $-3.25 \times 10^{-8}$ |
| 4 | 5 | $4.34 \times 10^{-2}$ | $4.30 \times 10^{-5}$ | $1.87 \times 10^{-6}$ | 2 | 3 | $-4.44 \times 10^{-3}$ | $3.28 \times 10^{-7}$ | $-1.46 \times 10^{-9}$ |
| 4 | 3 | $1.05 \times 10^{-2}$ | $6.75 \times 10^{-5}$ | $7.09 \times 10^{-7}$ | 3 | 2 | $-4.44 \times 10^{-3}$ | $3.20 \times 10^{-7}$ | $-1.42 \times 10^{-9}$ |

| 5'-15 DNA construct + 0.5 μM gp32 | | | | | | | | | |
| --- | --- | --- | --- | --- | --- | --- | --- | --- | --- |
| Positive terms |  |  |  |  | Negative terms |  |  |  |  |
| $i$ | $j$ | $\delta \bar{E}_i \delta \bar{E}_j$ | $w_{ij}$ | $w_{ij} \delta \bar{E}_i \delta \bar{E}_j$ | $i$ | $j$ | $\delta \bar{E}_i \delta \bar{E}_j$ | $w_{ij}$ | $w_{ij} \delta \bar{E}_i \delta \bar{E}_j$ |
| 1 | 1 | $1.67 \times 10^{-2}$ | $3.87 \times 10^{-1}$ | $6.48 \times 10^{-3}$ | 4 | 1 | $-3.50 \times 10^{-2}$ | $1.03 \times 10^{-2}$ | $-3.60 \times 10^{-4}$ |
| 4 | 4 | $7.32 \times 10^{-2}$ | $4.82 \times 10^{-2}$ | $3.53 \times 10^{-3}$ | 1 | 4 | $-3.50 \times 10^{-2}$ | $1.03 \times 10^{-2}$ | $-3.60 \times 10^{-4}$ |
| 3 | 3 | $8.26 \times 10^{-3}$ | $4.09 \times 10^{-1}$ | $3.38 \times 10^{-3}$ | 1 | 3 | $-1.18 \times 10^{-2}$ | $5.93 \times 10^{-5}$ | $-7.00 \times 10^{-7}$ |
| 5 | 5 | $1.37 \times 10^{-1}$ | $5.87 \times 10^{-3}$ | $8.07 \times 10^{-4}$ | 5 | 1 | $-4.79 \times 10^{-2}$ | $1.07 \times 10^{-5}$ | $-5.10 \times 10^{-7}$ |
| 2 | 2 | $8.59 \times 10^{-4}$ | $1.29 \times 10^{-1}$ | $1.11 \times 10^{-4}$ | 1 | 5 | $-4.79 \times 10^{-2}$ | $7.74 \times 10^{-5}$ | $-3.70 \times 10^{-7}$ |
| 5 | 4 | $1.00 \times 10^{-1}$ | $1.06 \times 10^{-4}$ | $1.07 \times 10^{-5}$ | 3 | 1 | $-1.18 \times 10^{-2}$ | $2.69 \times 10^{-5}$ | $-3.20 \times 10^{-7}$ |
| 4 | 5 | $1.00 \times 10^{-1}$ | $7.69 \times 10^{-5}$ | $7.72 \times 10^{-6}$ | 4 | 2 | $-7.93 \times 10^{-3}$ | $8.89 \times 10^{-7}$ | $-7.10 \times 10^{-9}$ |
| 3 | 5 | $3.37 \times 10^{-2}$ | $3.48 \times 10^{-5}$ | $1.17 \times 10^{-6}$ | 2 | 4 | $-7.93 \times 10^{-3}$ | $8.87 \times 10^{-7}$ | $-7.00 \times 10^{-9}$ |
| 2 | 1 | $3.79 \times 10^{-3}$ | $6.53 \times 10^{-5}$ | $2.47 \times 10^{-7}$ | 2 | 3 | $-2.66 \times 10^{-3}$ | $4.95 \times 10^{-9}$ | $-1.30 \times 10^{-11}$ |
| 1 | 2 | $3.79 \times 10^{-3}$ | $6.53 \times 10^{-5}$ | $2.47 \times 10^{-7}$ | 5 | 2 | $-1.09 \times 10^{-3}$ | $6.08 \times 10^{-10}$ | $-6.60 \times 10^{-12}$ |

| 3'-14 DNA construct + 0.5 μM gp32 | | | | | | | | | |
| --- | --- | --- | --- | --- | --- | --- | --- | --- | --- |
| Positive terms |  |  |  |  | Negative terms |  |  |  |  |
| $i$ | $j$ | $\delta \bar{E}_i \delta \bar{E}_j$ | $w_{ij}$ | $w_{ij} \delta \bar{E}_i \delta \bar{E}_j$ | $i$ | $j$ | $\delta \bar{E}_i \delta \bar{E}_j$ | $w_{ij}$ | $w_{ij} \delta \bar{E}_i \delta \bar{E}_j$ |
| 5 | 5 | $6.99 \times 10^{-2}$ | $5.83 \times 10^{-2}$ | $4.08 \times 10^{-3}$ | 4 | 1 | $-3.30 \times 10^{-2}$ | $1.30 \times 10^{-2}$ | $-4.30 \times 10^{-4}$ |
| 1 | 1 | $2.95 \times 10^{-2}$ | $1.26 \times 10^{-1}$ | $3.70 \times 10^{-3}$ | 1 | 4 | $-3.30 \times 10^{-2}$ | $1.29 \times 10^{-2}$ | $-4.30 \times 10^{-4}$ |
| 4 | 4 | $3.70 \times 10^{-2}$ | $4.36 \times 10^{-2}$ | $1.61 \times 10^{-3}$ | 1 | 3 | $-3.99 \times 10^{-3}$ | $3.65 \times 10^{-4}$ | $-1.50 \times 10^{-6}$ |
| 2 | 2 | $6.80 \times 10^{-3}$ | $1.93 \times 10^{-1}$ | $1.31 \times 10^{-3}$ | 5 | 1 | $-4.50 \times 10^{-2}$ | $2.45 \times 10^{-5}$ | $-1.10 \times 10^{-6}$ |
| 3 | 3 | $5.41 \times 10^{-4}$ | $5.47 \times 10^{-1}$ | $2.96 \times 10^{-4}$ | 3 | 1 | $-3.99 \times 10^{-3}$ | $2.29 \times 10^{-4}$ | $-9.10 \times 10^{-7}$ |
| 3 | 5 | $6.15 \times 10^{-3}$ | $2.80 \times 10^{-3}$ | $1.72 \times 10^{-5}$ | 1 | 5 | $-4.54 \times 10^{-2}$ | $7.45 \times 10^{-6}$ | $-3.40 \times 10^{-7}$ |
| 5 | 3 | $6.15 \times 10^{-3}$ | $2.66 \times 10^{-3}$ | $1.63 \times 10^{-5}$ | 4 | 2 | $-1.59 \times 10^{-2}$ | $4.28 \times 10^{-6}$ | $-6.80 \times 10^{-8}$ |
| 5 | 4 | $5.08 \times 10^{-2}$ | $1.70 \times 10^{-4}$ | $8.64 \times 10^{-6}$ | 2 | 4 | $-1.59 \times 10^{-2}$ | $4.24 \times 10^{-6}$ | $-6.70 \times 10^{-8}$ |
| 4 | 5 | $5.08 \times 10^{-2}$ | $4.67 \times 10^{-5}$ | $2.37 \times 10^{-6}$ | 2 | 3 | $-1.92 \times 10^{-3}$ | $1.15 \times 10^{-7}$ | $-2.20 \times 10^{-10}$ |
| 2 | 1 | $1.42 \times 10^{-2}$ | $8.18 \times 10^{-5}$ | $1.16 \times 10^{-6}$ | 3 | 2 | $-1.92 \times 10^{-3}$ | $7.18 \times 10^{-8}$ | $-1.40 \times 10^{-10}$ |

| 5'-14 DNA construct + 0.5 μM gp32 | | | | | | | | | |
| --- | --- | --- | --- | --- | --- | --- | --- | --- | --- |
| Positive terms |  |  |  |  | Negative terms |  |  |  |  |
| $i$ | $j$ | $\delta \bar{E}_i \delta \bar{E}_j$ | $w_{ij}$ | $w_{ij} \delta \bar{E}_i \delta \bar{E}_j$ | $i$ | $j$ | $\delta \bar{E}_i \delta \bar{E}_j$ | $w_{ij}$ | $w_{ij} \delta \bar{E}_i \delta \bar{E}_j$ |
| 1 | 1 | $1.54 \times 10^{-2}$ | $2.88 \times 10^{-1}$ | $4.42 \times 10^{-3}$ | 4 | 1 | $-3.36 \times 10^{-2}$ | $7.91 \times 10^{-3}$ | $-2.70 \times 10^{-4}$ |
| 4 | 4 | $7.33 \times 10^{-2}$ | $5.32 \times 10^{-2}$ | $3.90 \times 10^{-3}$ | 1 | 4 | $-3.36 \times 10^{-2}$ | $7.86 \times 10^{-3}$ | $-2.60 \times 10^{-4}$ |
| 3 | 3 | $8.08 \times 10^{-3}$ | $3.03 \times 10^{-1}$ | $2.45 \times 10^{-3}$ | 1 | 3 | $-1.11 \times 10^{-2}$ | $1.21 \times 10^{-4}$ | $-1.40 \times 10^{-6}$ |
| 5 | 5 | $1.38 \times 10^{-1}$ | $6.90 \times 10^{-3}$ | $9.49 \times 10^{-4}$ | 3 | 1 | $-1.11 \times 10^{-2}$ | $6.70 \times 10^{-5}$ | $-7.50 \times 10^{-7}$ |
| 2 | 2 | $8.48 \times 10^{-4}$ | $3.32 \times 10^{-1}$ | $2.82 \times 10^{-4}$ | 5 | 1 | $-4.60 \times 10^{-2}$ | $6.00 \times 10^{-6}$ | $-2.80 \times 10^{-7}$ |
| 5 | 4 | $1.00 \times 10^{-1}$ | $8.39 \times 10^{-5}$ | $8.43 \times 10^{-6}$ | 1 | 5 | $-4.60 \times 10^{-2}$ | $2.35 \times 10^{-6}$ | $-1.10 \times 10^{-7}$ |
| 4 | 5 | $1.00 \times 10^{-1}$ | $3.29 \times 10^{-5}$ | $3.30 \times 10^{-6}$ | 4 | 2 | $-7.88 \times 10^{-3}$ | $5.27 \times 10^{-6}$ | $-4.20 \times 10^{-8}$ |
| 3 | 5 | $3.33 \times 10^{-2}$ | $6.8 \times 10^{-5}$ | $2.27 \times 10^{-6}$ | 2 | 4 | $-7.88 \times 10^{-3}$ | $5.24 \times 10^{-6}$ | $-4.10 \times 10^{-8}$ |
| 2 | 1 | $3.61 \times 10^{-3}$ | $3.79 \times 10^{-4}$ | $1.37 \times 10^{-6}$ | 2 | 3 | $-2.62 \times 10^{-3}$ | $7.90 \times 10^{-8}$ | $-2.10 \times 10^{-10}$ |
| 1 | 2 | $3.61 \times 10^{-3}$ | $3.79 \times 10^{-4}$ | $1.37 \times 10^{-6}$ | 3 | 2 | $-2.62 \times 10^{-2}$ | $4.36 \times 10^{-8}$ | $-1.10 \times 10^{-10}$ |

**Table S8.** Decomposition of the largest pathway terms for the three-point TCFs [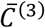 function, Eq. (8) of main text] at *τ*_1_ = *τ*_2_ = 1 ms.

| 3'-15 DNA construct + 0.1 μM gp32 | | | | | | | | | | | |
| --- | --- | --- | --- | --- | --- | --- | --- | --- | --- | --- | --- |
| Positive terms |  |  |  |  |  | Negative terms |  |  |  |  |  |
| $i$ | $j$ | $k$ | $\delta \bar{E}_i \delta \bar{E}_j \delta \bar{E}_k$ | $w_{ijk}$ | $w_{ijk} \delta \bar{E}_i \delta \bar{E}_j \delta \bar{E}_k$ | $i$ | $j$ | $k$ | $\delta \bar{E}_i \delta \bar{E}_j \delta \bar{E}_k$ | $w_{ijk}$ | $w_{ijk} \delta \bar{E}_i \delta \bar{E}_j \delta \bar{E}_k$ |
| 1 | 1 | 1 | $1.45 \times 10^{-3}$ | $4.97 \times 10^{-2}$ | $7.21 \times 10^{-5}$ | 3 | 3 | 3 | $-4.71 \times 10^{-3}$ | $4.05 \times 10^{-2}$ | $-1.91 \times 10^{-4}$ |
| 4 | 4 | 1 | $9.31 \times 10^{-3}$ | $7.39 \times 10^{-3}$ | $6.88 \times 10^{-5}$ | 1 | 4 | 1 | $-3.67 \times 10^{-3}$ | $1.92 \times 10^{-2}$ | $-7.05 \times 10^{-5}$ |
| 1 | 4 | 4 | $9.31 \times 10^{-3}$ | $7.37 \times 10^{-3}$ | $6.86 \times 10^{-5}$ | 4 | 1 | 1 | $-3.67 \times 10^{-3}$ | $1.92 \times 10^{-2}$ | $-7.04 \times 10^{-5}$ |
| 4 | 1 | 4 | $9.31 \times 10^{-3}$ | $7.36 \times 10^{-3}$ | $6.85 \times 10^{-5}$ | 1 | 1 | 4 | $-3.67 \times 10^{-3}$ | $1.91 \times 10^{-2}$ | $-7.02 \times 10^{-5}$ |
| 5 | 5 | 1 | $9.70 \times 10^{-3}$ | $3.73 \times 10^{-4}$ | $3.62 \times 10^{-6}$ | 4 | 4 | 4 | $-2.36 \times 10^{-2}$ | $2.84 \times 10^{-3}$ | $-6.70 \times 10^{-5}$ |
| 5 | 4 | 1 | $9.50 \times 10^{-3}$ | $2.84 \times 10^{-4}$ | $2.70 \times 10^{-6}$ | 5 | 5 | 5 | $-2.51 \times 10^{-2}$ | $6.06 \times 10^{-4}$ | $-1.52 \times 10^{-5}$ |
| 5 | 1 | 4 | $9.50 \times 10^{-3}$ | $2.48 \times 10^{-4}$ | $2.36 \times 10^{-6}$ | 5 | 5 | 4 | $-2.46 \times 10^{-2}$ | $1.63 \times 10^{-4}$ | $-4.00 \times 10^{-6}$ |
| 1 | 5 | 5 | $9.70 \times 10^{-3}$ | $1.26 \times 10^{-4}$ | $1.22 \times 10^{-6}$ | 5 | 4 | 4 | $-2.41 \times 10^{-2}$ | $1.09 \times 10^{-4}$ | $-2.62 \times 10^{-6}$ |
| 1 | 4 | 5 | $9.50 \times 10^{-3}$ | $9.52 \times 10^{-5}$ | $9.05 \times 10^{-7}$ | 5 | 1 | 1 | $-3.75 \times 10^{-3}$ | $6.46 \times 10^{-4}$ | $-2.42 \times 10^{-6}$ |
| 3 | 3 | 1 | $3.18 \times 10^{-3}$ | $2.59 \times 10^{-4}$ | $8.25 \times 10^{-7}$ | 4 | 5 | 5 | $-2.46 \times 10^{-2}$ | $5.48 \times 10^{-5}$ | $-1.35 \times 10^{-6}$ |

| 3'-15 DNA construct + 1.0 μM gp32 | | | | | | | | | | | |
| --- | --- | --- | --- | --- | --- | --- | --- | --- | --- | --- | --- |
| Positive terms |  |  |  |  |  | Negative terms |  |  |  |  |  |
| $i$ | $j$ | $k$ | $\delta \bar{E}_i \delta \bar{E}_j \delta \bar{E}_k$ | $w_{ijk}$ | $w_{ijk} \delta \bar{E}_i \delta \bar{E}_j \delta \bar{E}_k$ | $i$ | $j$ | $k$ | $\delta \bar{E}_i \delta \bar{E}_j \delta \bar{E}_k$ | $w_{ijk}$ | $w_{ijk} \delta \bar{E}_i \delta \bar{E}_j \delta \bar{E}_k$ |
| 2 | 2 | 2 | $7.79 \times 10^{-4}$ | $1.25 \times 10^{-1}$ | $9.73 \times 10^{-5}$ | 1 | 4 | 1 | $-3.82 \times 10^{-3}$ | $1.51 \times 10^{-2}$ | $-5.80 \times 10^{-5}$ |
| 1 | 1 | 1 | $4.60 \times 10^{-3}$ | $1.55 \times 10^{-2}$ | $7.11 \times 10^{-5}$ | 4 | 1 | 1 | $-3.82 \times 10^{-3}$ | $1.51 \times 10^{-2}$ | $-5.70 \times 10^{-5}$ |
| 4 | 4 | 1 | $3.17 \times 10^{-3}$ | $1.59 \times 10^{-2}$ | $5.03 \times 10^{-5}$ | 1 | 1 | 4 | $-3.82 \times 10^{-3}$ | $1.50 \times 10^{-2}$ | $-5.70 \times 10^{-5}$ |
| 1 | 4 | 4 | $3.17 \times 10^{-3}$ | $1.59 \times 10^{-2}$ | $5.03 \times 10^{-5}$ | 4 | 4 | 4 | $-2.63 \times 10^{-3}$ | $1.67 \times 10^{-2}$ | $-4.40 \times 10^{-5}$ |
| 4 | 1 | 4 | $3.17 \times 10^{-3}$ | $1.46 \times 10^{-2}$ | $4.64 \times 10^{-5}$ | 5 | 5 | 5 | $-7.30 \times 10^{-3}$ | $1.22 \times 10^{-3}$ | $-8.90 \times 10^{-6}$ |
| 2 | 2 | 1 | $1.41 \times 10^{-3}$ | $5.08 \times 10^{-3}$ | $7.15 \times 10^{-6}$ | 3 | 3 | 3 | $-8.20 \times 10^{-6}$ | $6.62 \times 10^{-1}$ | $-5.40 \times 10^{-6}$ |
| 1 | 2 | 2 | $1.40 \times 10^{-3}$ | $5.08 \times 10^{-3}$ | $7.15 \times 10^{-6}$ | 4 | 1 | 2 | $-2.11 \times 10^{-3}$ | $2.29 \times 10^{-3}$ | $-4.80 \times 10^{-6}$ |
| 2 | 1 | 1 | $2.54 \times 10^{-3}$ | $2.35 \times 10^{-3}$ | $5.99 \times 10^{-6}$ | 2 | 1 | 4 | $-2.11 \times 10^{-3}$ | $2.29 \times 10^{-3}$ | $-4.80 \times 10^{-6}$ |
| 1 | 1 | 2 | $2.54 \times 10^{-3}$ | $2.35 \times 10^{-3}$ | $5.99 \times 10^{-6}$ | 4 | 2 | 2 | $-1.17 \times 10^{-3}$ | $3.04 \times 10^{-3}$ | $-3.60 \times 10^{-6}$ |
| 5 | 4 | 1 | $4.45 \times 10^{-3}$ | $7.52 \times 10^{-4}$ | $3.35 \times 10^{-6}$ | 2 | 2 | 4 | $-1.17 \times 10^{-3}$ | $3.04 \times 10^{-3}$ | $-3.50 \times 10^{-6}$ |

| 3'-15 DNA construct + 0.5 μM gp32 | | | | | | | | | | | |
| --- | --- | --- | --- | --- | --- | --- | --- | --- | --- | --- | --- |
| Positive terms |  |  |  |  |  | Negative terms |  |  |  |  |  |
| $i$ | $j$ | $k$ | $\delta \bar{E}_i \delta \bar{E}_j \delta \bar{E}_k$ | $w_{ijk}$ | $w_{ijk} \delta \bar{E}_i \delta \bar{E}_j \delta \bar{E}_k$ | $i$ | $j$ | $k$ | $\delta \bar{E}_i \delta \bar{E}_j \delta \bar{E}_k$ | $w_{ijk}$ | $w_{ijk} \delta \bar{E}_i \delta \bar{E}_j \delta \bar{E}_k$ |
| 2 | 2 | 2 | $6.18 \times 10^{-4}$ | $3.72 \times 10^{-2}$ | $2.30 \times 10^{-4}$ | 4 | 4 | 4 | $-8.17 \times 10^{-3}$ | $6.34 \times 10^{-3}$ | $-5.20 \times 10^{-5}$ |
| 4 | 4 | 1 | $5.76 \times 10^{-3}$ | $8.08 \times 10^{-3}$ | $4.65 \times 10^{-5}$ | 1 | 4 | 1 | $-4.06 \times 10^{-3}$ | $1.02 \times 10^{-2}$ | $-4.20 \times 10^{-5}$ |
| 1 | 4 | 4 | $5.76 \times 10^{-3}$ | $8.05 \times 10^{-3}$ | $4.64 \times 10^{-5}$ | 4 | 1 | 1 | $-4.06 \times 10^{-3}$ | $9.71 \times 10^{-3}$ | $-3.90 \times 10^{-5}$ |
| 4 | 1 | 4 | $5.76 \times 10^{-3}$ | $7.61 \times 10^{-3}$ | $4.38 \times 10^{-5}$ | 1 | 1 | 4 | $-4.06 \times 10^{-3}$ | $9.68 \times 10^{-3}$ | $-3.90 \times 10^{-5}$ |
| 1 | 1 | 1 | $2.86 \times 10^{-3}$ | $1.24 \times 10^{-2}$ | $3.54 \times 10^{-5}$ | 3 | 3 | 3 | $-1.40 \times 10^{-4}$ | $2.55 \times 10^{-1}$ | $-3.60 \times 10^{-5}$ |
| 4 | 1 | 3 | $1.49 \times 10^{-3}$ | $8.79 \times 10^{-3}$ | $1.31 \times 10^{-5}$ | 3 | 3 | 5 | $-5.90 \times 10^{-4}$ | $2.69 \times 10^{-2}$ | $-1.60 \times 10^{-5}$ |
| 3 | 1 | 4 | $1.49 \times 10^{-3}$ | $8.67 \times 10^{-3}$ | $1.29 \times 10^{-5}$ | 5 | 3 | 3 | $-5.90 \times 10^{-4}$ | $2.66 \times 10^{-2}$ | $-1.60 \times 10^{-5}$ |
| 1 | 4 | 3 | $1.49 \times 10^{-3}$ | $7.86 \times 10^{-3}$ | $1.17 \times 10^{-5}$ | 3 | 5 | 3 | $-5.90 \times 10^{-4}$ | $2.46 \times 10^{-2}$ | $-1.40 \times 10^{-5}$ |
| 3 | 4 | 1 | $1.49 \times 10^{-3}$ | $7.84 \times 10^{-3}$ | $1.17 \times 10^{-5}$ | 3 | 5 | 5 | $-2.42 \times 10^{-3}$ | $5.88 \times 10^{-3}$ | $-1.40 \times 10^{-5}$ |
| 1 | 3 | 3 | $3.87 \times 10^{-4}$ | $2.47 \times 10^{-2}$ | $9.54 \times 10^{-6}$ | 5 | 5 | 3 | $-2.42 \times 10^{-3}$ | $5.81 \times 10^{-3}$ | $-1.40 \times 10^{-5}$ |

| 5'-15 DNA construct + 0.5 μM gp32 | | | | | | | | | | | |
| --- | --- | --- | --- | --- | --- | --- | --- | --- | --- | --- | --- |
| Positive terms |  |  |  |  |  | Negative terms |  |  |  |  |  |
| $i$ | $j$ | $k$ | $\delta \bar{E}_i \delta \bar{E}_j \delta \bar{E}_k$ | $w_{ijk}$ | $w_{ijk} \delta \bar{E}_i \delta \bar{E}_j \delta \bar{E}_k$ | $i$ | $j$ | $k$ | $\delta \bar{E}_i \delta \bar{E}_j \delta \bar{E}_k$ | $w_{ijk}$ | $w_{ijk} \delta \bar{E}_i \delta \bar{E}_j \delta \bar{E}_k$ |
| 1 | 1 | 1 | $2.16 \times 10^{-3}$ | $2.91 \times 10^{-1}$ | $6.29 \times 10^{-4}$ | 3 | 3 | 3 | $-7.50 \times 10^{-4}$ | $4.03 \times 10^{-1}$ | $-3.00 \times 10^{-4}$ |
| 4 | 4 | 1 | $9.48 \times 10^{-3}$ | $6.27 \times 10^{-3}$ | $5.94 \times 10^{-5}$ | 4 | 1 | 1 | $-4.53 \times 10^{-3}$ | $4.28 \times 10^{-2}$ | $-1.90 \times 10^{-4}$ |
| 4 | 1 | 4 | $9.48 \times 10^{-3}$ | $6.27 \times 10^{-3}$ | $5.94 \times 10^{-5}$ | 1 | 1 | 4 | $-4.53 \times 10^{-3}$ | $4.26 \times 10^{-2}$ | $-1.90 \times 10^{-4}$ |
| 1 | 4 | 4 | $9.48 \times 10^{-3}$ | $6.25 \times 10^{-3}$ | $5.92 \times 10^{-5}$ | 1 | 4 | 1 | $-4.53 \times 10^{-3}$ | $4.26 \times 10^{-2}$ | $-1.90 \times 10^{-4}$ |
| 5 | 5 | 1 | $1.78 \times 10^{-3}$ | $1.12 \times 10^{-3}$ | $1.99 \times 10^{-5}$ | 5 | 5 | 5 | $-5.09 \times 10^{-2}$ | $8.18 \times 10^{-4}$ | $-4.20 \times 10^{-5}$ |
| 1 | 5 | 5 | $1.78 \times 10^{-3}$ | $8.07 \times 10^{-4}$ | $1.43 \times 10^{-5}$ | 4 | 4 | 4 | $-1.98 \times 10^{-2}$ | $9.21 \times 10^{-4}$ | $-1.80 \times 10^{-5}$ |
| 5 | 4 | 1 | $1.30 \times 10^{-2}$ | $5.58 \times 10^{-4}$ | $7.24 \times 10^{-6}$ | 5 | 1 | 1 | $-6.20 \times 10^{-3}$ | $2.59 \times 10^{-6}$ | $-1.60 \times 10^{-5}$ |
| 1 | 4 | 5 | $1.30 \times 10^{-2}$ | $4.02 \times 10^{-4}$ | $5.22 \times 10^{-6}$ | 1 | 1 | 5 | $-6.20 \times 10^{-3}$ | $1.87 \times 10^{-3}$ | $-1.20 \times 10^{-5}$ |
| 5 | 1 | 4 | $1.30 \times 10^{-2}$ | $3.79 \times 10^{-4}$ | $4.92 \times 10^{-6}$ | 5 | 5 | 4 | $-3.72 \times 10^{-2}$ | $2.41 \times 10^{-4}$ | $-9.00 \times 10^{-6}$ |
| 4 | 1 | 5 | $1.30 \times 10^{-2}$ | $2.75 \times 10^{-4}$ | $3.56 \times 10^{-6}$ | 1 | 5 | 1 | $-6.20 \times 10^{-3}$ | $1.10 \times 10^{-3}$ | $-6.80 \times 10^{-6}$ |

| 3'-14 DNA construct + 0.5 μM gp32 | | | | | | | | | | | |
| --- | --- | --- | --- | --- | --- | --- | --- | --- | --- | --- | --- |
| Positive terms |  |  |  |  |  | Negative terms |  |  |  |  |  |
| $i$ | $j$ | $k$ | $\delta \bar{E}_i \delta \bar{E}_j \delta \bar{E}_k$ | $w_{ijk}$ | $w_{ijk} \delta \bar{E}_i \delta \bar{E}_j \delta \bar{E}_k$ | $i$ | $j$ | $k$ | $\delta \bar{E}_i \delta \bar{E}_j \delta \bar{E}_k$ | $w_{ijk}$ | $w_{ijk} \delta \bar{E}_i \delta \bar{E}_j \delta \bar{E}_k$ |
| 1 | 1 | 1 | $5.06 \times 10^{-3}$ | $5.42 \times 10^{-2}$ | $2.74 \times 10^{-4}$ | 4 | 1 | 1 | $-5.67 \times 10^{-3}$ | $2.21 \times 10^{-2}$ | $-1.30 \times 10^{-4}$ |
| 2 | 2 | 2 | $5.60 \times 10^{-4}$ | $1.84 \times 10^{-1}$ | $1.03 \times 10^{-4}$ | 1 | 4 | 1 | $-5.67 \times 10^{-3}$ | $2.21 \times 10^{-2}$ | $-1.30 \times 10^{-4}$ |
| 4 | 4 | 1 | $6.35 \times 10^{-3}$ | $9.02 \times 10^{-3}$ | $5.73 \times 10^{-5}$ | 1 | 1 | 4 | $-5.67 \times 10^{-3}$ | $2.19 \times 10^{-2}$ | $-1.20 \times 10^{-4}$ |
| 4 | 1 | 4 | $6.35 \times 10^{-3}$ | $8.96 \times 10^{-3}$ | $5.69 \times 10^{-5}$ | 5 | 5 | 5 | $-1.85 \times 10^{-2}$ | $1.50 \times 10^{-3}$ | $-2.80 \times 10^{-5}$ |
| 1 | 4 | 4 | $6.35 \times 10^{-3}$ | $8.94 \times 10^{-3}$ | $5.68 \times 10^{-5}$ | 4 | 4 | 4 | $-7.12 \times 10^{-3}$ | $3.65 \times 10^{-3}$ | $-2.60 \times 10^{-5}$ |
| 5 | 4 | 1 | $8.73 \times 10^{-3}$ | $8.33 \times 10^{-4}$ | $7.27 \times 10^{-6}$ | 5 | 1 | 1 | $-7.79 \times 10^{-3}$ | $1.95 \times 10^{-3}$ | $-1.50 \times 10^{-5}$ |
| 5 | 1 | 4 | $8.73 \times 10^{-3}$ | $7.88 \times 10^{-4}$ | $6.88 \times 10^{-6}$ | 3 | 5 | 5 | $-1.63 \times 10^{-3}$ | $7.74 \times 10^{-3}$ | $-1.30 \times 10^{-5}$ |
| 5 | 5 | 1 | $1.20 \times 10^{-2}$ | $4.90 \times 10^{-4}$ | $5.88 \times 10^{-6}$ | 5 | 5 | 3 | $-1.63 \times 10^{-3}$ | $7.38 \times 10^{-3}$ | $-1.20 \times 10^{-5}$ |
| 2 | 1 | 1 | $2.43 \times 10^{-3}$ | $1.81 \times 10^{-3}$ | $4.39 \times 10^{-6}$ | 1 | 1 | 5 | $-7.79 \times 10^{-3}$ | $9.73 \times 10^{-4}$ | $-7.60 \times 10^{-6}$ |
| 1 | 1 | 2 | $2.43 \times 10^{-3}$ | $1.81 \times 10^{-3}$ | $4.39 \times 10^{-6}$ | 5 | 3 | 5 | $-1.63 \times 10^{-3}$ | $4.23 \times 10^{-3}$ | $-6.90 \times 10^{-6}$ |

| 5'-14 DNA construct + 0.5 μM gp32 | | | | | | | | | | | |
| --- | --- | --- | --- | --- | --- | --- | --- | --- | --- | --- | --- |
| Positive terms |  |  |  |  |  | Negative terms |  |  |  |  |  |
| $i$ | $j$ | $k$ | $\delta \bar{E}_i \delta \bar{E}_j \delta \bar{E}_k$ | $w_{ijk}$ | $w_{ijk} \delta \bar{E}_i \delta \bar{E}_j \delta \bar{E}_k$ | $i$ | $j$ | $k$ | $\delta \bar{E}_i \delta \bar{E}_j \delta \bar{E}_k$ | $w_{ijk}$ | $w_{ijk} \delta \bar{E}_i \delta \bar{E}_j \delta \bar{E}_k$ |
| 1 | 1 | 1 | $1.90 \times 10^{-3}$ | $1.75 \times 10^{-1}$ | $3.34 \times 10^{-4}$ | 3 | 3 | 3 | $-7.30 \times 10^{-4}$ | $2.90 \times 10^{-1}$ | $-2.10 \times 10^{-4}$ |
| 4 | 4 | 1 | $9.09 \times 10^{-3}$ | $7.60 \times 10^{-3}$ | $6.91 \times 10^{-5}$ | 1 | 4 | 1 | $-4.16 \times 10^{-3}$ | $3.65 \times 10^{-2}$ | $-1.50 \times 10^{-4}$ |
| 1 | 4 | 4 | $9.09 \times 10^{-3}$ | $7.55 \times 10^{-3}$ | $6.86 \times 10^{-5}$ | 4 | 1 | 1 | $-4.16 \times 10^{-3}$ | $3.65 \times 10^{-2}$ | $-1.50 \times 10^{-4}$ |
| 4 | 1 | 4 | $9.09 \times 10^{-3}$ | $7.53 \times 10^{-3}$ | $6.84 \times 10^{-5}$ | 1 | 1 | 4 | $-4.16 \times 10^{-3}$ | $3.62 \times 10^{-2}$ | $-1.50 \times 10^{-4}$ |
| 5 | 5 | 1 | $1.70 \times 10^{-2}$ | $1.11 \times 10^{-3}$ | $1.90 \times 10^{-5}$ | 5 | 5 | 5 | $-5.10 \times 10^{-2}$ | $1.60 \times 10^{-3}$ | $-8.20 \times 10^{-5}$ |
| 1 | 5 | 5 | $1.70 \times 10^{-3}$ | $4.44 \times 10^{-4}$ | $7.56 \times 10^{-6}$ | 4 | 4 | 4 | $-1.99 \times 10^{-2}$ | $1.57 \times 10^{-3}$ | $-3.10 \times 10^{-5}$ |
| 2 | 2 | 2 | $2.47 \times 10^{-5}$ | $2.97 \times 10^{-1}$ | $7.34 \times 10^{-6}$ | 3 | 5 | 5 | $-1.24 \times 10^{-2}$ | $1.15 \times 10^{-3}$ | $-1.40 \times 10^{-5}$ |
| 5 | 4 | 1 | $1.24 \times 10^{-2}$ | $5.88 \times 10^{-4}$ | $7.32 \times 10^{-6}$ | 5 | 5 | 4 | $-3.72 \times 10^{-2}$ | $3.62 \times 10^{-4}$ | $-1.30 \times 10^{-5}$ |
| 2 | 1 | 1 | $4.47 \times 10^{-4}$ | $1.17 \times 10^{-2}$ | $5.25 \times 10^{-6}$ | 5 | 1 | 1 | $-5.69 \times 10^{-3}$ | $1.79 \times 10^{-3}$ | $-1.00 \times 10^{-5}$ |
| 1 | 1 | 2 | $4.47 \times 10^{-4}$ | $1.17 \times 10^{-2}$ | $5.25 \times 10^{-6}$ | 3 | 3 | 5 | $-3.00 \times 10^{-3}$ | $2.34 \times 10^{-3}$ | $-7.00 \times 10^{-6}$ |

**Figure S4.**
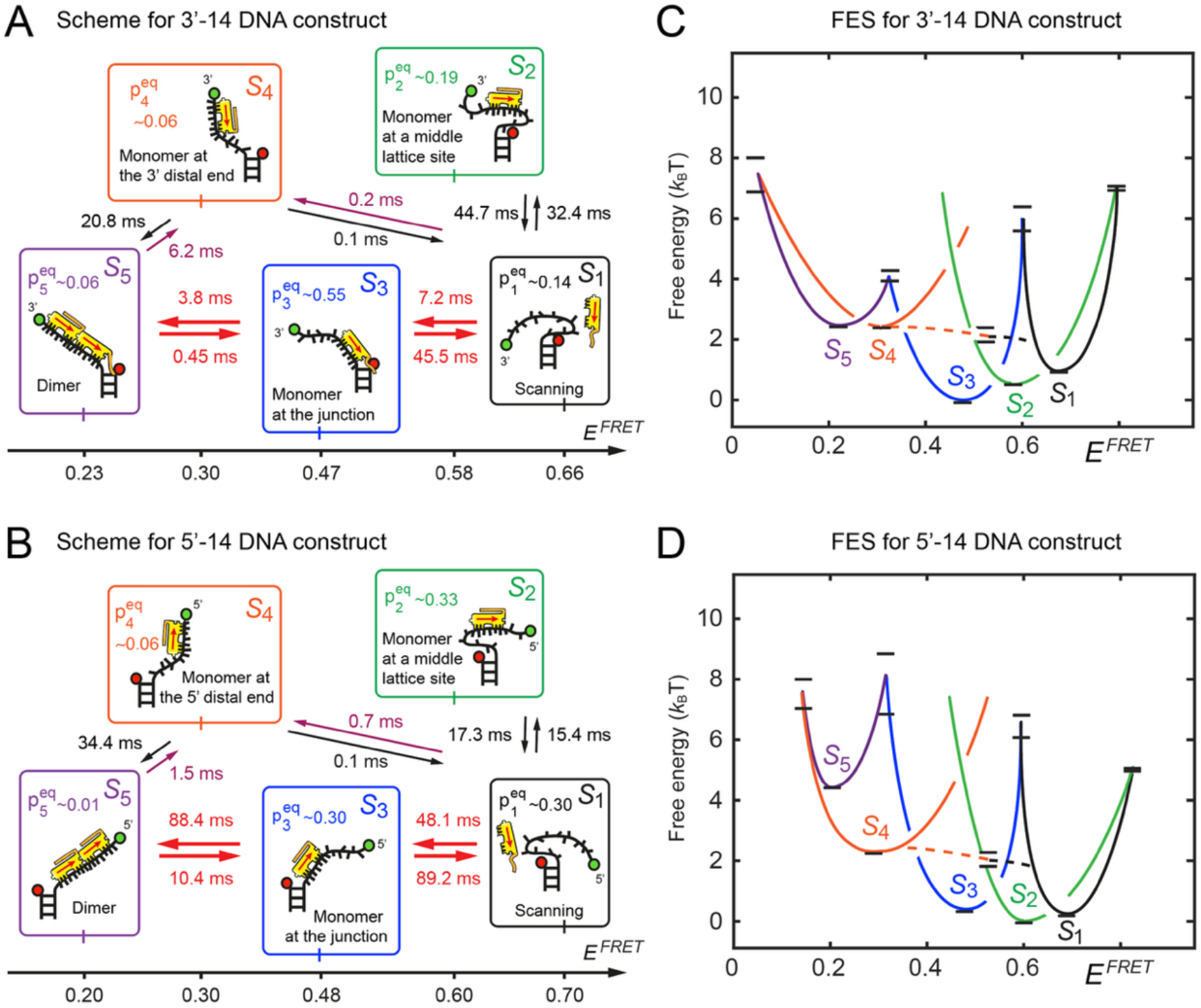
Kinetic network schemes and FESs for (***A, C***) the 3’-oligo(dT)_14_-dsDNA construct (3’-14) and (***B, D***) the 5’-oligo(dT)_14_-dsDNA construct (5’-14) in the presence of [gp32] = 0.5 *μ*M in aqueous buffer with 100 mM NaCl and 6 mM MgCl_2_. Optimized kinetic and equilibrium parameters are listed, as in Fig. 4 of the main text. The structural assignments for macrostates *S*_1_ – *S*_5_ are depicted schematically indicating the orientation of the CTD relative to the ss-dsDNA junction and the polarity of the oligo(dT)_14_ overhanging tail.

## Notes

### Competing Interest Statement

The authors have declared no competing interest.

